# Aβ oligomers trigger necroptosis-mediated neurodegeneration via microglia activation in Alzheimer’s disease

**DOI:** 10.1101/2021.08.27.457960

**Authors:** Natalia Salvadores, Inés Moreno-Gonzalez, Nazaret Gamez Ruiz, Gabriel Quiroz, Laura Vegas, Marcela Escandón, Sebastian Jimenez, Javier Vitorica, Antonia Gutierrez, Claudio Soto, Felipe A. Court

**Affiliations:** Center for Integrative Biology, Faculty of Sciences, Universidad Mayor, Santiago, Chile; FONDAP Geroscience Center for Brain Health and Metabolism, Santiago, Chile; Department of Cell Biology, Facultad de Ciencias, Universidad de Malaga, IBIMA, Malaga, Spain; Networking Research Center on Neurodegenerative Diseases (CIBERNED), Madrid, Spain; Mitchell Center for Alzheimer’s Disease, Department of Neurology, McGovern Medical School, University of Texas Medical School at Houston, Houston, TX; Dpto. Bioquimica y Biologia Molecular, Facultad de Farmacia, Universidad de Sevilla, 41012 Sevilla, Spain; Instituto de Biomedicina de Sevilla (IBiS)-Hospital Universitario Virgen del Rocío/CSIC/Universidad de Sevilla, 41013 Sevilla, Spain; Buck Institute for Research on Ageing, Novato, San Francisco

**Keywords:** Alzheimer’s disease, Amyloid-β oligomers, necroptosis, microglia, neurodegeneration, neuroprotection

## Abstract

Alzheimer’s disease (AD) is a major adult-onset neurodegenerative condition with no available treatment. Compelling reports point amyloid-β (Aβ) as the main etiologic agent that triggers AD. Although there is extensive evidence of detrimental crosstalk between Aβ and microglia that contributes to neuroinflammation in AD, the exact mechanism leading to neuron death remains unknown. Using postmortem human AD brain tissue, we show that Aβ pathology is associated with the necroptosis effector pMLKL. Moreover, we found that the burden of Aβo correlates with the expression of key markers of necroptosis activation. Additionally, inhibition of necroptosis by pharmacological or genetic means, reduce neurodegeneration and memory impairment triggered by Aβo in mice. Since microglial activation is emerging as a central driver for AD pathogenesis, we then tested the contribution of microglia to the mechanism of Aβo-mediated necroptosis activation in neurons. Using an *in vitro* model, we show that conditioned medium from Aβo-stimulated microglia elicited necroptosis in neurons through activation of TNF-α signaling, triggering extensive neurodegeneration. Notably, necroptosis inhibition provided significant neuronal protection. Together, these findings suggest that Aβo-mediated microglia stimulation in AD contributes to necroptosis activation in neurons and neurodegeneration. As necroptosis is a druggable degenerative mechanism, our findings might have important therapeutic implications to prevent the progression of AD.

## Introduction

Alzheimer’s disease (AD) represents the most frequent brain disorder in humans, affecting mainly the elderly population. AD is characterized by progressive neurodegeneration leading to impaired cognition, memory loss, and dementia (1). The neuropathological hallmarks of AD include the accumulation of intraneuronal deposits of hyperphosphorylated tau (pTau) protein and intracellular and extracellular aggregates of amyloid-β (Aβ), which are associated with dystrophic neurites and activated microglia (2). Increasing studies have demonstrated that soluble oligomeric Aβ structures contribute importantly to neurodegeneration in AD and correlate with cognitive alterations (2–7).

Familial AD, which represents less than 5% of the cases, is caused by mutations in the amyloid precursor protein (APP) gene or mutations in the presenilin 1 or 2 genes, leading to increased deposition of Aβ (1, 2). The most common form of the disease is sporadic AD with unknown origin, where aging is the most important risk factor. Accumulating evidence has attributed a causative role to Aβ in the etiopathogenesis of AD (2, 8). Neuroinflammation is a critical component of the pathogenesis of Aβ, involving abnormal activation of glial cells and the production of various proinflammatory cytokines (9). Following the first description of activated microglial cells in close contact with amyloid plaques in AD brains, extensive research focusing on the role of microglia in AD pathogenesis has been conducted (10–14). Upon interaction with Aβ, activated microglia initiate an innate immune response that leads to sustained production and secretion of pro-inflammatory mediators including interleukin (IL)-6, IL-1β, and tumor necrosis factor-α (TNF-α), contributing to neuronal damage and cognitive decline (15–19). Interestingly, mutations in the Triggering Receptor Expressed on Myeloid cells 2 (TREM2), which is expressed in microglia, increase the risk of AD by about 3-to 5-fold (20, 21). TREM2 is involved in microglial signaling and plays an important role in Aβ phagocytosis as well as suppressing cytokine production and neuroinflammation (22–24). These studies support an important contribution of inflammatory responses in the mechanism of Aβ-induced neurodegeneration.

Necroptosis is a mechanism of programmed cell death that participates in the pathogenesis of several diseases (25). Necroptosis is activated in postmortem human AD brains and the expression of necroptotic markers correlates with the disease stage and progression at the cognitive level (26, 27). Nevertheless, the mechanism that triggers necroptosis in AD remains to be established. Different stimuli induce necroptosis activation, including TNF ligand family members (28), interferons (29), and toll-like receptor 3 stimulation (30). Stimulation of TNF type 1 receptor (TNFR1) leads to the recruitment of receptor-interacting protein kinase 1 (RIPK1) (31, 32), which triggers the activation of either caspase-8 or RIPK3, leading to apoptosis or necroptosis, respectively (33). Following a stimulus such as TNF-α ligation, and in the absence of caspase-8 activity, the interaction of RIPK1 and RIPK3 induces the formation of the necrosome complex and RIPK3 phosphorylation, a key event in necroptosis activation (34–36). RIPK3 activation leads to the recruitment and phosphorylation of the pseudokinase mixed lineage kinase domain-like protein (MLKL) within the necrosome complex. Upon activation, MLKL is oligomerized and executes necroptosis leading to cell death (37, 38). Interestingly, recent studies suggest that necroptosis may operate as a relevant degenerative pathway in models of amyotrophic lateral sclerosis (ALS) (39), multiple sclerosis (MS) (40), and Parkinson’s disease (PD) (41).

Here, we investigated the role of Aβ pathology in the mechanism underlying necroptosis activation in AD. We found that Aβ oligomers (Aβo) are associated with key necroptosis drivers in the brains of AD patients and that targeting this death mechanism reduces Aβo-related neurodegeneration and memory impairment in mice. Furthermore, we report that *in vitro*, Aβo elicit necroptosis-mediated neurodegeneration via microglia activation through the engagement of TNF-α signaling. This study sheds light on the role of necroptosis in the crosstalk between glial and neuronal cells in AD and provides evidence indicating that necroptosis constitutes a potential therapeutic target for the disease.

## Methods

### Human samples

Histological analyses: Hippocampal brain samples were obtained from the Neurological Tissue Bank of IDIBELL-Hospital of Bellvitge (Barcelona, Spain) fixed in formaldehyde and embedded in paraffin. Research on human samples was performed following the code of ethics of the World Medical Association (Declaration of Helsinki). Samples were manipulated following the universal precautions for working with human samples and as directed by the supplier biobank and local ethic committees following Spanish legislation.

Molecular and biochemical analyses: autopsy specimens from the medial temporal lobe (hippocampal/parahippocampal regions) were obtained from the tissue bank Fundación CIEN (BT-CIEN; Centro de Investigación de Enfermedades Neurologicas; Madrid, Spain) and from the Neurological Tissue Bank of IDIBELL-Hospital of Bellvitge (Barcelona, Spain). The utilization of postmortem human samples was approved by the corresponding biobank ethics committees and by the “‘‘Comite de Etica de la Investigacion (CEI), Hospital Virgen del Rocio”‘‘, Seville, Spain. We considered the age-matched Braak II individuals as controls in our experiments. Only Braak V–VI cases were clinically classified as demented (AD) patients.

### Mice

MLKL knockout mice were kindly provided by Dr. Douglas Green (St. Jude Children’s Research Hospital, Memphis, TN, USA) and have been described previously (42); male (4-6 months old) mice were used, wild-type littermates were used as controls. For the GSK’872 experiment, C57BL/6 mice were used and were obtained from Jackson Laboratory (Bar Harbor, ME, USA), male (13-14 months old) mice were used. Animals were housed in groups of a maximum of five in individually ventilated cages under standard conditions (22 °C, 12 h light-dark cycle) receiving food and water ad libitum. All animal manipulations were carried out in accordance with standard regulations and approved by the Animal Care and Use Scientific Ethics Committee of the Mayor University.

### Intracerebroventricular injection

Mice were anesthetized with isoflurane and placed in a stereotaxic frame (David Kopf Instruments, USA). A single cerebroventricular injection was performed in both lateral ventricles according to the following stereotaxic coordinates: -1.0 ± 0.06 mm posterior to bregma, 1.8 ± 0.1 mm lateral to the sagittal suture, and 2.4 mm in depth, as previously described (43). Five μl of a solution of Aβo (100 μM) or vehicle (PBS) were injected in each ventricle at a rate of 0.5 μl/min.

### GSK’872 administration

GSK’872 (Tocris Bioscience) was dissolved in DMSO to a final concentration of 100 mM (stock). The working solution was prepared by diluting the stock solution with DMSO and then sterile PBS at room temperature to 0.3 mM (20% DMSO). The vehicle consisted of a solution of 20% DMSO in PBS. GSK’872 or vehicle were administrated via intraperitoneal injection (1 mg/Kg) weekly for three weeks, starting the same day of the surgery.

### Behavioral assessment

Two weeks after surgery, cognitive impairment was tested using the Morris Water Maze as previously described (44). In this assessment, using visual cues for orientation, animals learn to swim to a hidden platform under the water. During the training, mice are placed into the pool and allowed to explore for 1 min. If the animals do not find the platform, they are guided to it. This training procedure was performed 4 times a day for four consecutive days per animal. On day five, the platform was removed from the pool, and the time that the animals spent in the target quadrant was measured. At the end of the experiments, animals were euthanized by an overdose of anesthesia, and all efforts were made to minimize suffering. Then, animals were transcardially perfused with 50 mL of PBS.

### Preparation of synthetic oligomers

Aβo were prepared as previously described (45). Briefly, lyophilized recombinant Aβ42 peptide (rPeptide, A-1163) was re-suspended to 5 mM in DMSO and then further diluted to 100 μM in cold Neurobasal medium for the *in vitro* experiments or in sterile PBS for the *in vivo* experiments. The peptide was incubated at 4 °C for 24 h to generate oligomers, which were characterized by electron microscopy.

### Total RNA extraction, retrotranscription, and quantitative real-time PCR

Total RNA was extracted from human samples using TriPure Isolation Reagent (Roche). RNA integrity (RIN) was determined by RNA Nano 6000 (Agilent). No differences between the Braak groups were observed (RIN: 6.95 ± 1.4). RNA was quantified using NanoDrop 2000 spectrophotometer (Thermo Fischer). Retrotranscription (RT) (4 μg of total RNA) was performed with the High-Capacity cDNA Archive Kit (Applied Biosystems). For quantitative real-time PCR (qPCR), 40 ng of cDNA were mixed with 2× Taqman Universal Master Mix (Applied Biosystems) and 20× Taqman Gene Expression assay probes (*Ripk1*, Hs01041869_m1; *Ripk3*, Hs00179132_m1; *Mlkl*, Hs04188505_m1; Applied Biosystems). qPCR reactions were done using an ABI Prism 7900HT (Applied Biosystems). The cDNA levels were determined using GAPDH and beta-actin. We observed a highly significant linear correlation between the cycle threshold (Ct) of both genes (beta-actin vs GAPDH, *r* = 0.912, *F*(1,46) = 157.73, *p* < 0.0001). Thus, normalization using either beta-actin (Hs99999903_1, Applied Biosystems) or GAPDH (Hs03929097_g1, Applied Biosystems) produced identical results. Routinely, we used GAPDH as housekeeper. Results were expressed using the comparative double-delta Ct method (2-ΔΔCt). ΔCt values represent GAPDH normalized expression levels. ΔΔCt was calculated using Braak II.

### Preparation of soluble S1 fractions

Soluble S1 fractions from human samples were prepared as described (46). Briefly, human tissue was homogenized (Dounce homogenizer) in TBS (20 mM Tris–HCl, 140 mM NaCl, pH 7.5) containing protease and phosphatase inhibitors (Roche). Homogenates were ultracentrifuged (4 °C for 60 min) at 100,000×*g* (Optima MAX Preparative Ultracentrifuge, Beckman Coulter). Supernatants (S1 fractions), were aliquoted and stored at −80 °C.

### Tissue processing for histopathology

For the study of human samples, deparaffinized 10-μm-thick sections were used. After auto-fluorescence blocking (Millipore), sections were subjected to antigen retrieval by heating at 80°C for 30 min in 10 mM citrate buffer pH 6.0. Samples were blocked with 5% BSA diluted in PBS Tx-100 (0.2%). For double staining, samples were incubated with anti-pMLKL (1:200, Sabbiotech) for 48 h and stained with Thioflavin-S (0.025% in PBS) for 8 min. Signal was detected incubating for 1.5 h with the corresponding Alexa secondary antibody (1:500, Invitrogen). For double immunostaining with anti-pMLKL (1:200, Sabbiotech) and anti-MAP2 (1:5000, Thermo Fisher) or anti-Iba-1 antibodies (1:1000, Wako), samples were incubated overnight at 4 °C. Then, sections were incubated with the corresponding Alexa secondary antibodies (1:1000). For triple staining, samples were incubated for 10 min in Amylo-Glo® RTD (Biosensis), washed, and consecutively incubated with anti-pMLKL and anti-Iba-1 antibodies. Signal was detected as described above. Finally, samples were treated with an antifade mounting medium (VECTASHIELD). Fluorescent microscopy images were taken in a Leica SP8 confocal laser microscope coupled to a computer with TCS NT software (Leica).

For the histological study on mice, following perfusion, the whole brain was quickly removed, the right hemisphere was fixed with 4% PFA and then incubated in 30% sucrose before for OCT-embedding. Brains were sagittally cut into 20-μm-thick serial slices using a cryostat. Tissue sections were rinsed with TBS and then mounted onto superfrost plus slides. Antigen retrieval was performed by submerging slices in 10 mM citrate buffer pH 6.0 at 100 °C for 20 min. Samples were blocked with 5% BSA and then incubated with the antibody solution (anti-pMLKL 1:200, Abcam; anti-Iba1 1:700, Wako), overnight at 4 °C. Finally, antigens were visualized by incubating with the corresponding Alexa secondary antibodies (1:500, Thermo Fisher). Fluoro-Jade C staining: the FJC ready-to-dilute staining kit (Biosensis) was used. Briefly, sections were immersed in a basic alcohol solution consisting of 1% sodium hydroxide in 80% ethanol for 5 min, followed by 2 min in 70% ethanol and 2 min in distilled water. Then, sections were incubated for 10 min in a solution of 0.06% potassium permanganate. Samples were rinsed in distilled water and stained for 10 min with the FJ-C solution. After rinsing, the air-dried slides were cleared in xylene for 1 min and then coverslipped. Images were obtained using a Leica DMi8 microscope.

### Image analysis

All analyzes were carried out using Image J software (v1.42q). Colocalization analysis: The Coloc2 plugin was used to calculate the Manders Correlation values for intensities above Costes threshold. These values were used to obtain the ratio of pixels from one channel that colocalizes with total pixels in the second channel. The percentage of stained area was determined in thresholded images, by quantifying the signal above threshold divided by the total image area. Total stained area in a specific ROI was determined in thresholded images, by quantifying the signal above threshold in that ROI divided by the area of the ROI.

### Cell culture

Hippocampal neuronal cultures: Primary neuronal cultures were prepared from embryonic Sprague-Dawley rats (E18). Briefly, meninges-free hippocampi were isolated, trypsinized, and plated onto poly-L-lysine (0.05 mg/ml)-coated glass coverslips on tissue culture wells (150 cells/mm^2^). Neurons were grown in Neurobasal media with L-glutamine and B27 supplements to consistently provide neuronal cultures >95% pure. Cultures were treated with 1 μM AraC to inhibit glial cell proliferation. For all experiments, neurons were treated on *in vitro* day 7.

Microglial cell culture: Microglial cells were cultured from postnatal day 3 Sprague-Dawley rat brains. Briefly, meninges-free cortices were dissected, and cells were mechanically dissociated. Cells were plated onto tissue culture flasks in DMEM media containing 10% heat-inactivated fetal bovine serum. Culture medium was replaced on *in vitro* days 1 and 5. After 10 days, the cultures were shaken for 2 h at 180 rpm on a rotary shaker to remove microglia.

Co-cultures: Hippocampal and microglial cell cultures were performed as described above. Microglial cells were added to hippocampal neuronal cultures at a ratio of 1:1, at *in vitro* day 7, 4 h before Aβo treatments.

### Preparation of conditioned medium

Conditioned medium was prepared as previously described (47). Microglial cells (180 cells/mm^2^) were plated onto immobilized Aβo (7.3 pmol/ mm^2^) to prevent their subsequent collection in the conditioned medium. To immobilize the oligomers, tissue culture dishes were coated with nitrocellulose that was obtained by dissolving a piece of membrane in methanol. Oligomers were distributed on the dish and allowed to dry for 10 min. Then, the dish was washed twice and blocked with 5% BCA in DMEM. After 48 h of stimulation, the microglial conditioned medium was collected and stored at – 80 °C. To prepare the control medium, the same process was performed except that oligomers were not added. Additionally, a negative control was prepared by collecting medium from dishes containing immobilized Aβo alone.

### Immunofluorescence

Neurons were fixed in 4% PFA at room temperature for 20 min. Cells were then blocked using 5% fish gelatin, 0,1% triton-X-100 for 2 h. Then, cells were incubated with the primary antibody solution (anti-acetylated tubulin 1:1000, Sigma-Aldrich; anti-Iba1 1:000, Wako) over-night at 4 °C. After washing with PBS, cells were incubated with the secondary antibody solution (goat anti-mouse alexa 488 or 546 1:500, Thermo Fisher) for 1 h at room temperature. Cells were washed and coverslipped with Mowiol mounting medium (Sigma).

### Protein extraction, quantification, and co-immunoprecipitation

Briefly, cells were collected with a cell scraper and centrifuged at 5000 x g for 5 min. The pellet was re-suspended in RIPA buffer (50 mM Tris pH 7.5, 150 mM NaCl, 1% NP-40, 0.5% sodium deoxycholate, 0.1% SDS, 100 *μ*g/ml PMSF) with phosphatase and protease inhibitor cocktail and stored at - 80 °C. Protein levels were quantified with Pierce BCA Protein Assay Kit (Thermo Scientific). Immunoprecipitation experiments were performed by incubating 100 μg of protein with 2.5 μg of anti-RIPK1 (BD Transduction Laboratories) with rotation at 4 °C for 48 h. Then, 50 μl of protein G magnetic beads (Biorad) were added to each sample and incubated with rotation at 4 °C for 3 h. Following magnetic separation, beads were mixed with loading buffer and boiled at 90 °C for 5 min. After the elution step, samples were analyzed by western blotting as described.

### Western blot

Proteins were fractionated by electrophoresis using 10% sodium dodecyl sulphate (SDS) polyacrylamide gels, electroblotted into PVDF membranes (Hybond-P, GE Healthcare), and blocked with 5% BCA in TBS. Membranes were then incubated with the different antibodies (anti-RIPK3, abcam ab16090; anti-pMLKL, abcam ab196436, and anti-HSP90, Santa Cruz sc7947) overnight at 4 °C, followed by incubation with a horseradish peroxidase-conjugated antibody. The immunoreactive bands were visualized using ECL (Invitrogen). The images were obtained and analyzed with ChemiDoc™ Touch Imaging System (Bio-Rad). For the immunoprecipitation experiments, detection of the IgG was done following the detection of pMLKL, using membranes that were stripped with a solution containing glycine 0,2 M, SDS 0,1%, Tween 20 0,1 %, pH 2,2 (2 × 5 min incubation). After confirming that no bands could be visualized with ECL, membranes were blocked and incubated with a horseradish peroxidase-conjugated anti-mouse antibody.

### Dot blot

Dot blots were done as described previously (48). Briefly, 1 μg of protein from the different S1 fractions was first treated with PureProteome Protein G Magnetics Beads (Millipore) for contaminant IgG depletion and then directly applied to dry nitrocellulose, in a final volume of 2 μl. Then, blots were blocked for 1 h and incubated overnight at 4 °C with anti-Amyloid fibrils (OC, 1:5000, Merck-Millipore), anti-prefibrillar oligomers (A11, 1:5000 InvitroGen), anti pMLKL (1:1000, abcam) or anti-GAPDH (1:10000, Cell Signalling) antibodies. Blots were visualized using the Pierce ECL 2 Western Blotting Substrate detection method (0.5 pg, lower limit sensitivity; Thermo Scientific). The images were obtained and analyzed with ChemiDoc™ Touch Imaging System (Bio-Rad). Data were always normalized by the specific signal observed in the Braak II group.

### Statistical analyses

All experiments were analyzed in a blinded manner. To analyze differences in the data obtained from molecular and biochemical analyses of human samples, data were compared by Kruskal-Wallis test followed by Dunn posthoc test. For the association analyses, linear regression was done using the Spearman rho-test. Data from all other studies were statistically analyzed using Student’s *t-test* (for two groups comparisons) or one-way ANOVA (for more than two groups) followed by Bonferroni’s post hoc test. For the association analyses, linear regression was done using Pearson correlation. Data are shown as mean ± SEM. All analyses were performed using Graph Pad Prism 5.0 software. Differences were considered significant with p < 0.05.

## Results

### Aβ pathology is associated with necroptosis activation in human AD brains

Although evidence for necroptosis activation has been reported in the brain of patients with AD (26, 27), the possible association between Aβ pathology and necroptosis activation has not been established. Therefore, we studied the association between Aβ and phosphorylated MLKL at Ser358 (pMLKL), which defines the activation of a canonical necroptotic process, in brain samples derived from the hippocampus of sporadic AD, mild cognitive impairment (MCI), and control (CTRL) cases (clinical data of the subjects used in the histopathological study are listed in Table 1). The histopathological assessment revealed low pMLKL immunoreactivity in the hippocampus of control samples (Fig. 1A). Of the four controls analyzed, three were Aβ plaque-free and did not present immunoreactivity for pMLKL (see CTRL1). The control sample that exhibited Aβ pathology presented a low extent of Aβ deposits, and some of them were associated with pMLKL (CTRL2, see arrows). Brain samples from AD cases showed considerable levels of Aβ plaques, and most of them depicted some degree (moderate to high) of positive pMLKL immunoreactivity. As shown in the merged images, pMLKL signal was observed within the Aβ plaques, and also surrounding the deposits. Remarkably, these findings were also observed in brain samples derived from patients with MCI, which is considered a precursor for AD (49). The association between Aβ plaques and pMLKL was additionally studied using the specific probe for amyloid plaques, Amylo-Glo (Figure 1B). The pMLKL immunoreactivity was validated using DAB staining (Supplementary Fig. 1). These data demonstrate a spatial correlation between Aβ pathology and the necroptosis effector in AD, which is consistent with the fact that Aβ plaques are often surrounded by dystrophic neurites and associated with synaptic loss (50).

**Table 1.** Clinical data of cases investigated by histological analysis of postmortem brain tissue. MMSE: Mini-mental state examination, CTRL: Control, MCI: Mild cognitive impairment, AD: Alzheimer’s disease, M: Male, F: Female, N/A: Not available.

| Condition | Gender | Age | Braak Stage | MMSE |
| --- | --- | --- | --- | --- |
| CTRL | M | 80 | I | 30 |
| CTRL | F | >89 | II | 30 |
| CTRL | N/A | N/A | II | N/A |
| CTRL | N/A | N/A | II | N/A |
| MCI | N/A | N/A | N/A | N/A |
| MCI | N/A | N/A | N/A | N/A |
| MCI | N/A | N/A | N/A | N/A |
| AD | M | 65 | III | N/A |
| AD | M | 84 | III | N/A |
| AD | F | 79 | IV | N/A |
| AD | M | 75 | VI | 0 |
| AD | F | 76 | VI | 1 |
| AD | F | >89 | VI | 6 |
| AD | M | 84 | VI | N/A |
| AD | M | 81 | VI | N/A |
| AD | F | 56 | VI | N/A |
| AD | F | 66 | VI | N/A |
| AD | F | 76 | VI | N/A |
| AD | F | 86 | VI | N/A |

**Figure 1.**
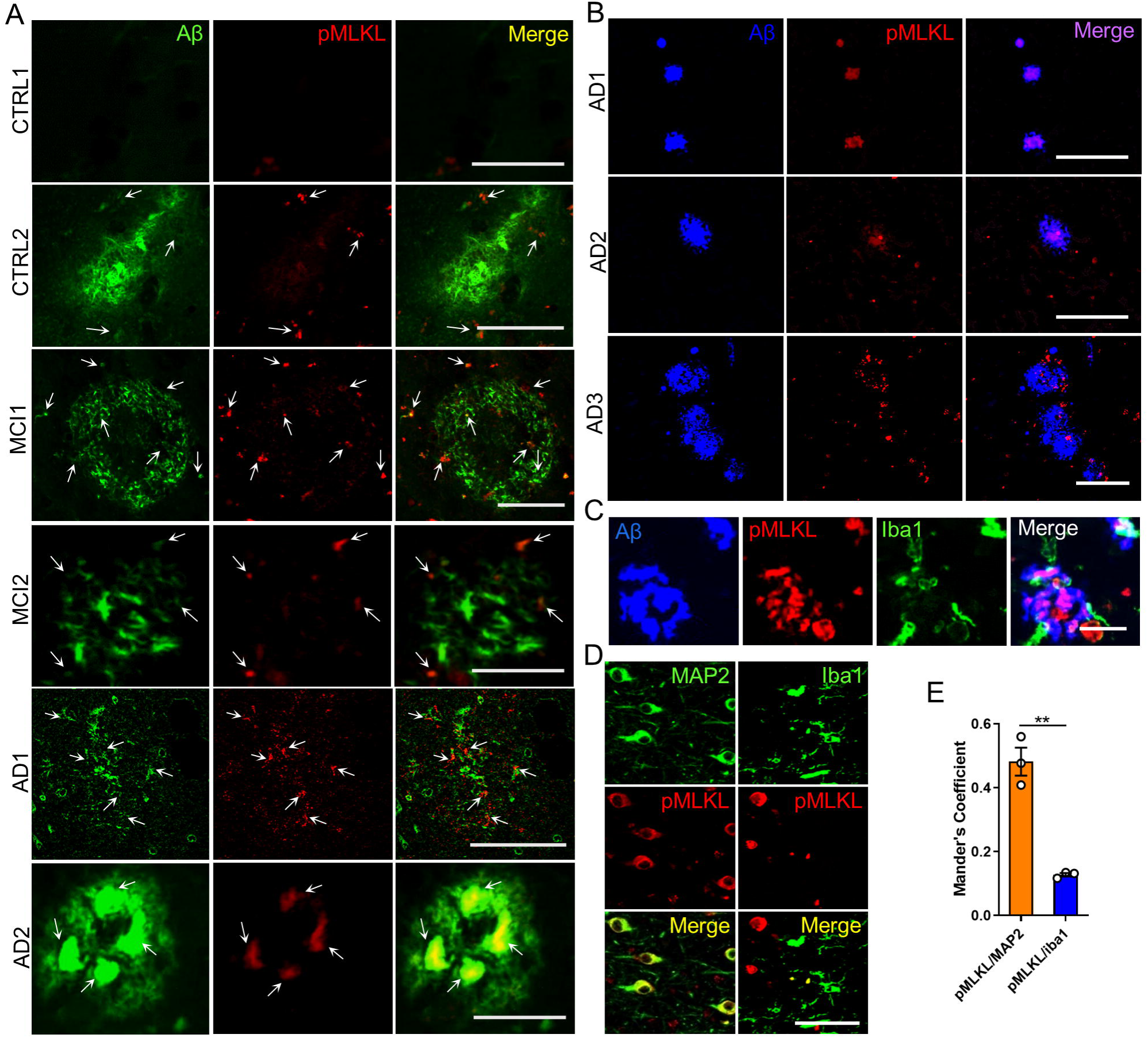
Aβ deposits colocalize with pMLKL in AD brains. **(A)** Representative pictures of the localization of pMLKL in relation to the thioflavin S-positive Aβ deposits in hippocampal brain areas of AD (n = 12), MCI, (n = 3), and CTRL (n = 4) patients (scale bar, CTRL1 25 μm; CTRL2 25 μm; MCI1 50 μm; MCI2 25 μm; AD1 100 μm; and AD2 25 μm). **(B)** Representative micrographs of the localization of pMLKL in relation to the Amylo-Glo-positive Aβ deposits in hippocampal brain areas of AD (n = 3) patients (scale bar, 100 μm). **(C)** Representative images of hippocampal areas of human AD brains (n = 3), stained with the Amylo-Glo Aβ marker (blue), and co-labeled with the indicated antibodies (scale bar, 25 μm). **(D**,**E)** Colocalization analyses between MAP2 (green) and pMLKL (red, left panel), and between Iba1 (green) and pMLKL (red, right panel) in hippocampal areas of human AD brains (n = 3) was done by determining the Manders coefficient (scale bar, 100 μm). Data are presented as mean ± S.E.M. Data in E were analyzed by Student’s *t-test*. **p < 0.01.

Microglial cells are commonly observed around amyloid deposits (11) (Fig. 1C). Since previous studies in other degenerative conditions have shown necroptosis activation in microglial cells (51, 52), we then tested whether pMLKL immunoreactivity localizes to neurons or microglia in AD. To this aim, we conducted colocalization analyses between pMLKL and MAP2 or Iba1, respectively, using AD brain tissue (Fig. 1D,E). The results revealed that most of the pMLKL immunoreactivity colocalized with MAP2. This result suggests that necroptosis activation occurs mainly in neurons of the AD brain.

Several lines of investigation indicate that Aβo oligomers (Aβo) contribute importantly to neurodegeneration in AD (2, 3). Indeed, previous studies have demonstrated that soluble oligomeric Aβ correlates better with cognitive deficits compared to plaques, which can actually sequester oligomers, acting as reservoirs from which oligomers can diffuse, reaching cellular targets in their vicinity (4–7). Thus, we next moved forward to determine the relationship between the burden of Aβo and necroptosis activation in AD brains. To this end, we evaluated the association between *Ripk1, Ripk3*, and *Mlkl* mRNA levels and the levels of soluble Aβo in postmortem brain samples of AD patients at different stages of the disease (clinical data of the subjects used in this study are summarized in Table 2). Samples were divided into three groups: samples derived from Braak II, non-demented individuals, which did not accumulate Aβ in the hippocampus; samples from Braak III-IV patients, which presented Aβ accumulation; and samples from Braak V-VI patients, which presented significantly higher levels of Aβ compared to the latter group (Figure 2A,B). In agreement with previously reported data (26), we observed that the expression levels of *Ripk1* and *Mlkl*, but not those of *Ripk3*, are elevated in the brains of individuals with higher Braak stages (Fig. 2C). The levels of oligomeric Aβ aggregates were assessed by dot blot using OC and A11 antibodies to detect soluble fibrillar and prefibrillar Aβo, respectively (Fig. 2D). The dynamic range and the coefficient of variation of the dot-blot assays are shown in Supplementary Fig.2. Quantitative analysis revealed that the extent of both types of structures was significantly higher in the brains of AD patients compared to healthy controls (Fig. 2E). We found a significant and positive correlation between Aβo load and the levels of *Ripk1* but not *Ripk3* or *MLKL* mRNA (Fig. 2F,G). Since MLKL activation is post-translationally regulated by phosphorylation, we then measured the association of pMLKL levels with the load of Aβ oligomers in AD brain samples. To achieve this, pMLKL levels were determined by dot blot (Fig. 2H). Quantitative measurement showed significantly increased pMLKL levels in AD subjects compared to controls (Fig. 2I). Notably, the levels of pMLKL correlate significantly and positively with the burden of Aβo (Fig. 2J,K). These results suggest a link between oligomeric Aβ structures and activation of the necroptotic machinery.

**Table 2.** Demographics of the individuals investigated by molecular and biochemical analyses of postmortem brain tissue.

| Braak Stage | Gender (Male, %) | Gender (Female, %) | Age (Years, mean age ± SD) | Postmortem delay (mean hours ± SD) |
| --- | --- | --- | --- | --- |
| II (n=10) | 61.54 | 38.46 | 78 ± 8.5 | 7 ± 4 |
| III-IV (n=9) | 44.44 | 55.56 | 80 ± 11 | 6 ± 5 |
| V-VI (n=10) | 38.89 | 61.11 | 79 ± 10 | 8 ± 4 |

**Figure 2.**
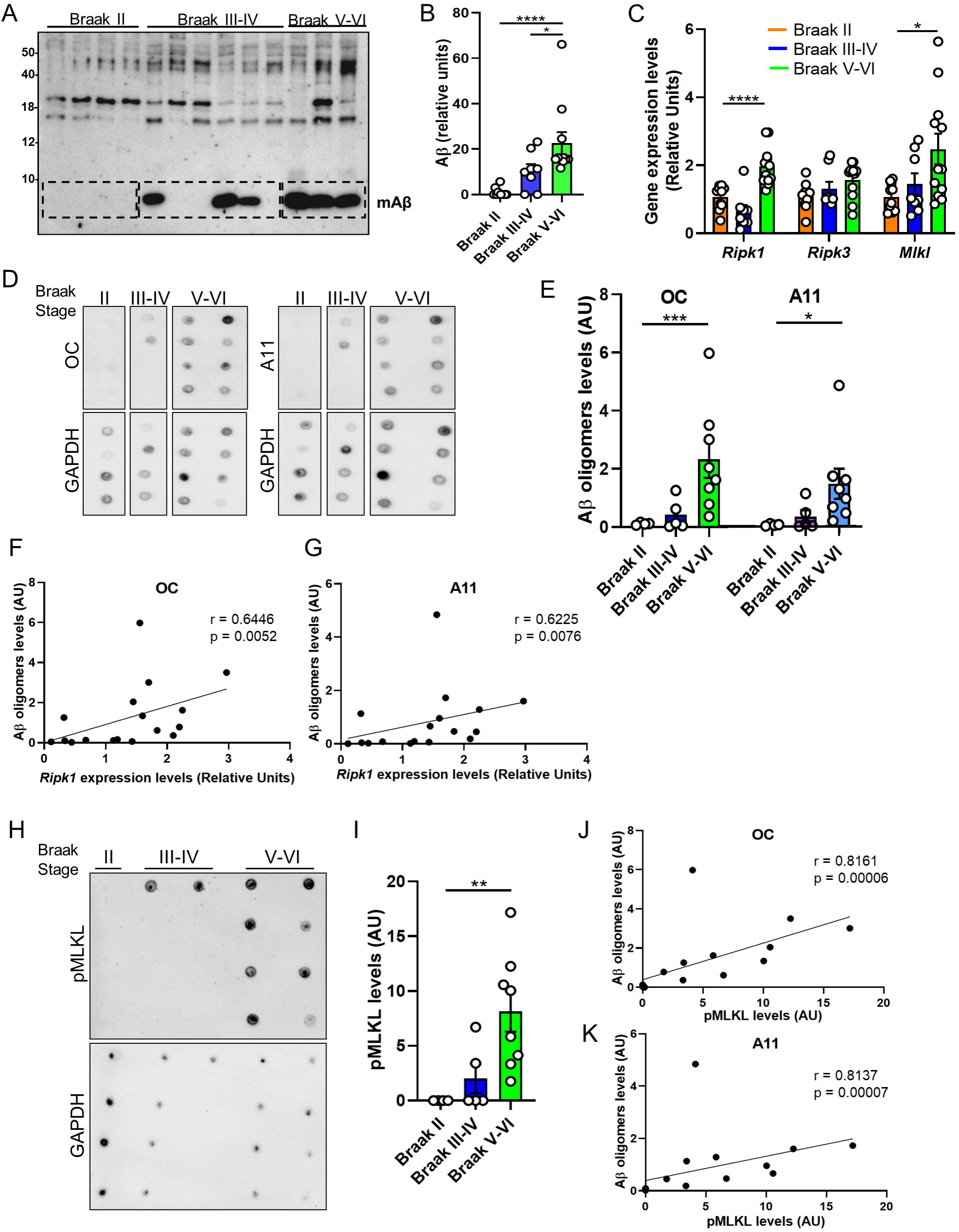
Activation of RIPK1 and MLKL correlate with soluble Aβ oligomer burden in AD brains. **(A)** Western blot analysis of total protein extracts from hippocampal samples using a mix of 6E10 and 82E1 antibodies was performed to determine the Aβ content. **(B)** The specific band was quantified by densitometric analysis, represented in the graphs. **(C)** The graph represents the expression levels of *Ripk1, Ripk3*, and *Mlkl* in human brain samples from patients at different AD stages (Braak stage II (n = 10), III-IV (n = 9), V-VI (n = 10)). **(D)** Dot blots of soluble proteins extracted from postmortem brains of individuals with Braak II (n = 4), Braak III-IV (n = 5), and Braak V-VI (n = 8), probed with the indicated antibodies. **(E)** Blots were quantified by densitometric analysis and expressed relative to GAPDH protein, represented in the graph. **(F**,**G)** Association analyses of oligomeric Aβ load and mRNA levels were done by linear regression. **(H)** Dot blots of protein extracts from postmortem brains of individuals with Braak II (n = 4), Braak III-IV (n = 5), and Braak V-VI (n = 8), probed with the indicated antibodies. **(I)** Blots were quantified by densitometric analysis and expressed relative to GAPDH protein, represented in the graph. **(J**,**K)** Association analyses of oligomeric Aβ load and pMLKL levels were done by linear regression. Data are presented as mean ± S.E.M. Data in **B, C, E**, and **I** were analyzed by Kruskal-Wallis test followed by Dunn post-hoc test. Data in **F, G, J** and **K** were analyzed by linear regression using Spearman rho-test. *p < 0.05; **p < 0.01; ***p < 0.001; ****p < 0.0001.

Together, these data provide the first evidence that Aβ pathology is associated with necroptosis activation in AD.

### Targeting necroptosis ameliorates neurodegeneration and memory impairment induced by Aβo in mice

Given the role of Aβ aggregates as major pathological mediators of both familial and sporadic AD and considering that Aβo are generally regarded as the most pathogenic form of Aβ, animal models based on intracerebral Aβo inoculation have been established as useful tools to study AD pathogenesis (3). To assess whether Aβo can induce activation of the necroptosis pathway, we conducted an acute intracerebroventricular injection of synthetic Aβo to wild-type mice. As expected, Aβo injection triggered significant microgliosis compared to control mice (Fig. 3A,B). Moreover, Aβo treated mice underwent neurodegeneration in the hippocampus, as demonstrated by an increase in non-phosphorylated neurofilaments (Fig. 3C,D). Staining for pMLKL demonstrated significant necroptosis activation in Aβo injected mice compared to vehicle-injected ones (Fig. 3E,F). To validate the phospho-specific MLKL antibody, we compared the immunoreactivity of brain sections from wild-type and MLKL knock-out mice, treated with Aβo (Supplementary Fig. 3A,B). In addition, to assess the cell specificity of the pMLKL immunostaining, we performed double labeling using antibodies to pMLKL and MAP2 or CD11b to observe neurons and microglia, respectively. As shown in Supplementary Fig. 3C,D, most of the pMLKL signal observed in the hippocampus was associated with neurons. Notably, pMLKL expression positively correlated with neurodegeneration determined by Fluoro-Jade C staining (Supplementary Fig. 4) and negatively correlated with cognitive performance evaluated using the Morris water maze (MWM) behavioral test (Fig. 3G,H).

**Figure 3.**
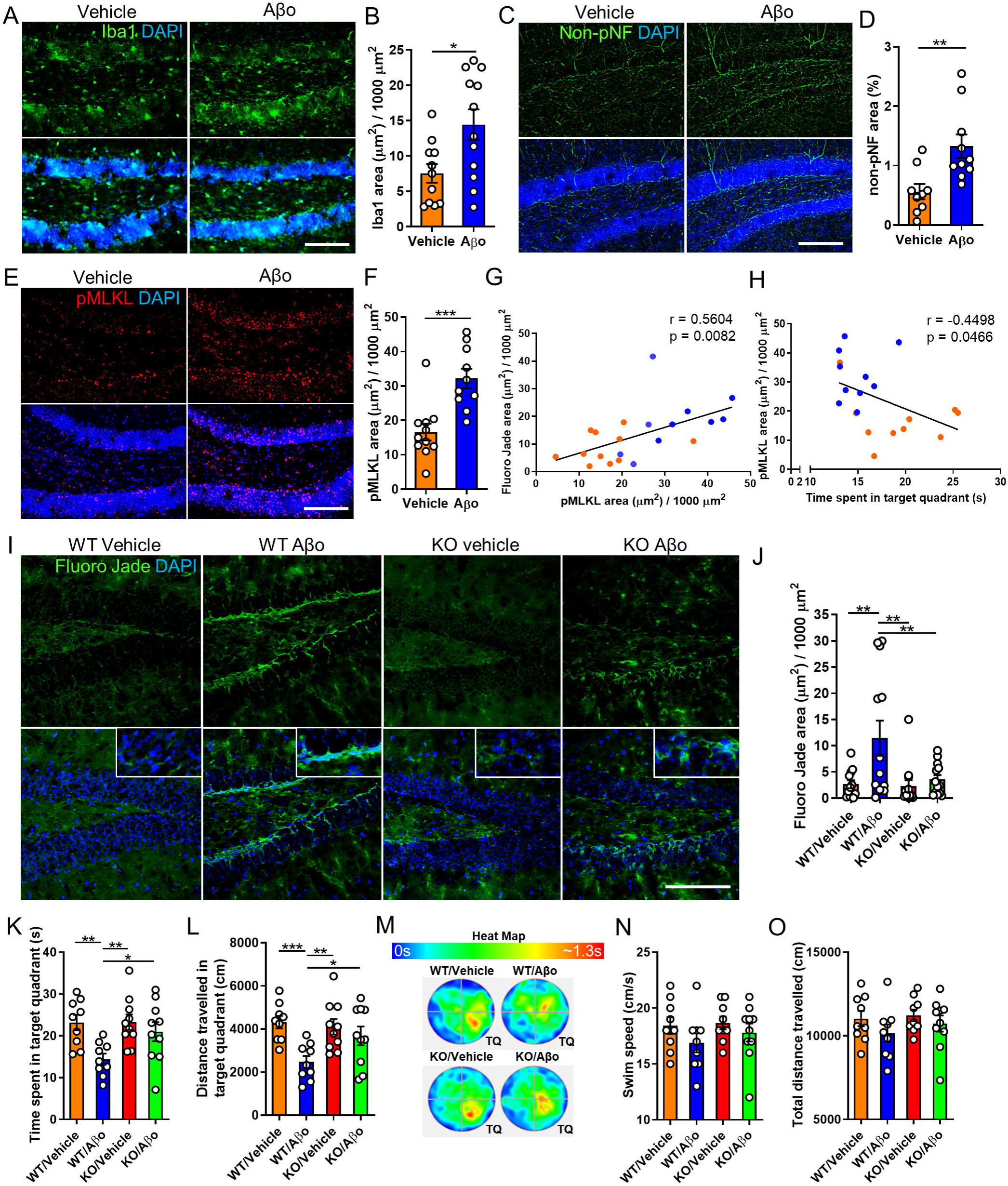
MLKL ablation ameliorates neurodegeneration and memory loss induced by Aβo in mice. **(A**,**B)** Representative micrographs and quantitative analysis of brain sections from wild-type mice subjected to intracerebroventricular infusion of Aβo or vehicle, immunostained with an Iba1 specific antibody (scale bar, 150 μm). **(C**,**D)** Representative images and quantitative analysis of brain sections immunolabeled with an anti-non-phosphorylated neurofilament antibody (scale bar, 150 μm). **(E**,**F)** Representative images and quantitative analysis of brain sections immunolabeled with an anti-pMLKL antibody (scale bar, 150 μm). Association analyses of **(G)** pMLKL and Fluoro-Jade C stained area and **(H)** pMLKL stained area and cognitive performance, were done by linear regression. **(I**,**J)** Representative micrographs and quantitative measurement of brain sections from mice, treated as indicated in the images, stained with Fluoro-Jade C (scale bar, 150 μm). n = 13 per group, 4 sections per brain, one image of the hippocampus per section was analyzed. **(K**,**L**,**N**,**O)** The graphs represent quantitative analyzes of the indicated parameters, obtained from the MWM test. **(M)** The heat map shows a graphic representation of the time that mice, treated as indicated in the image, spent in the target quadrant (TQ) during the MWM test. Data are presented as mean ± S.E.M. The data in **B, D, and F** were analyzed by Student’s *t-test*. Data in **G** and **H** were analyzed by linear regression using Pearson correlation. Data in **J**-**L, N**, and **O** were analyzed by one-way ANOVA followed by Bonferroni post-test. *p < 0.05; **p < 0.01; ***p < 0.001.

To evaluate the potential protective effect of MLKL inhibition against Aβo-induced neurodegeneration and memory alterations, we used genetically modified mice lacking *Mlkl*. To test the effects of targeting MLKL in the progression of AD pathogenesis, we evaluated neurodegeneration in the brain of these mice by histopathological assessment using Fluoro-Jade C staining. As shown in Fig. 3I,J, Aβo administration elicited significant neurodegeneration compared to control mice, evidenced by increased Fluoro-Jade C stained area in the hippocampus. Notably, ablation of *Mlkl* expression decreased the levels of Aβo-related neurodegeneration. To determine whether targeting MLKL influences the microgliosis observed following Aβo treatments, immunostaining was done using Iba1. Although similar levels of microglial activation were observed following Aβo infusion, in transgenic and wild-type mice, no statistically significant differences were found between controls and Aβo-treated mice in the knock-out group (Supplementary Fig. 5). In order to study the possible consequences of inhibiting MLKL on the behavioral impairment observed in this AD model, we evaluated the memory capacity of mice using the MWM test. In agreement with previous reports (53, 54) Aβo-injected mice exhibited altered memory performance in the MWM test, as this group of animals spent less time in the target quadrant compared to control mice (Fig. 3K). In sharp contrast, *Mlkl* deletion significantly inhibited the cognitive alteration observed in this AD model as the time spent in the target quadrant by MLKL^-/-^ mice treated with Aβo was significantly longer compared to MLKL^+/+^ mice injected with the oligomers (Fig. 3K). The protection provided by *Mlkl* deletion was also reflected when measuring the distance that mice traveled in the target quadrant during the test (Fig. 3L). The time that mice spent in the target quadrant during the test is graphically represented in the heat map (Fig. 3M). Importantly, swim speed and total traveled distance were not modified by Aβo treatment (Fig. 3N,O), suggesting this model does not cause locomotor or visual damage.

To increase the translational potential of these findings, we next investigated the possible protective effect of targeting necroptosis by pharmacological inhibition in this AD model. To this end, we treated mice with the RIPK3 inhibitor GSK’872. Histopathological exploration of brain pMLKL levels revealed that intraperitoneal administration of GSK’872 significantly decreased necroptosis activation induced by Aβo (Fig. 4A,B). Moreover, evaluation of neurodegeneration using Fluoro-Jade C showed that GSK’872 significantly attenuated Aβo-related neurodegeneration to levels comparable to those detected in control mice (Fig. 4C,D). To determine whether targeting RIPK3 influences the microgliosis triggered by Aβo treatment, immunostaining was done using Iba1. As shown in Supplementary Fig. 6, *Ripk3* ablation did not alter the microglial activation induced by Aβo, suggesting neuronal necroptosis engagement occurs downstream microglial activation. To evaluate the functional consequences of inhibiting RIPK3, cognitive performance was tested using the MWM. As expected, and similarly to the result described above, mice subjected to Aβo infusion underwent significant memory impairment. Remarkably, the results of the test revealed that treatment with GSK’872 provided significant protection against Aβo-induced memory loss, as mice challenged with Aβo and treated with the drug, behaved similarly to control animals (Fig. 4E,F). The time that mice spent in the target quadrant during the test is graphically represented in the heat map (Fig. 4G). The results shown in Fig. 4H,I suggest that Aβo treatment did not cause locomotor or visual damage in these mice. Noteworthy, a negative correlation between neurodegeneration assessed by Fluoro-Jade C and cognitive performance was observed in these mice (Fig. 4J).

**Figure 4.**
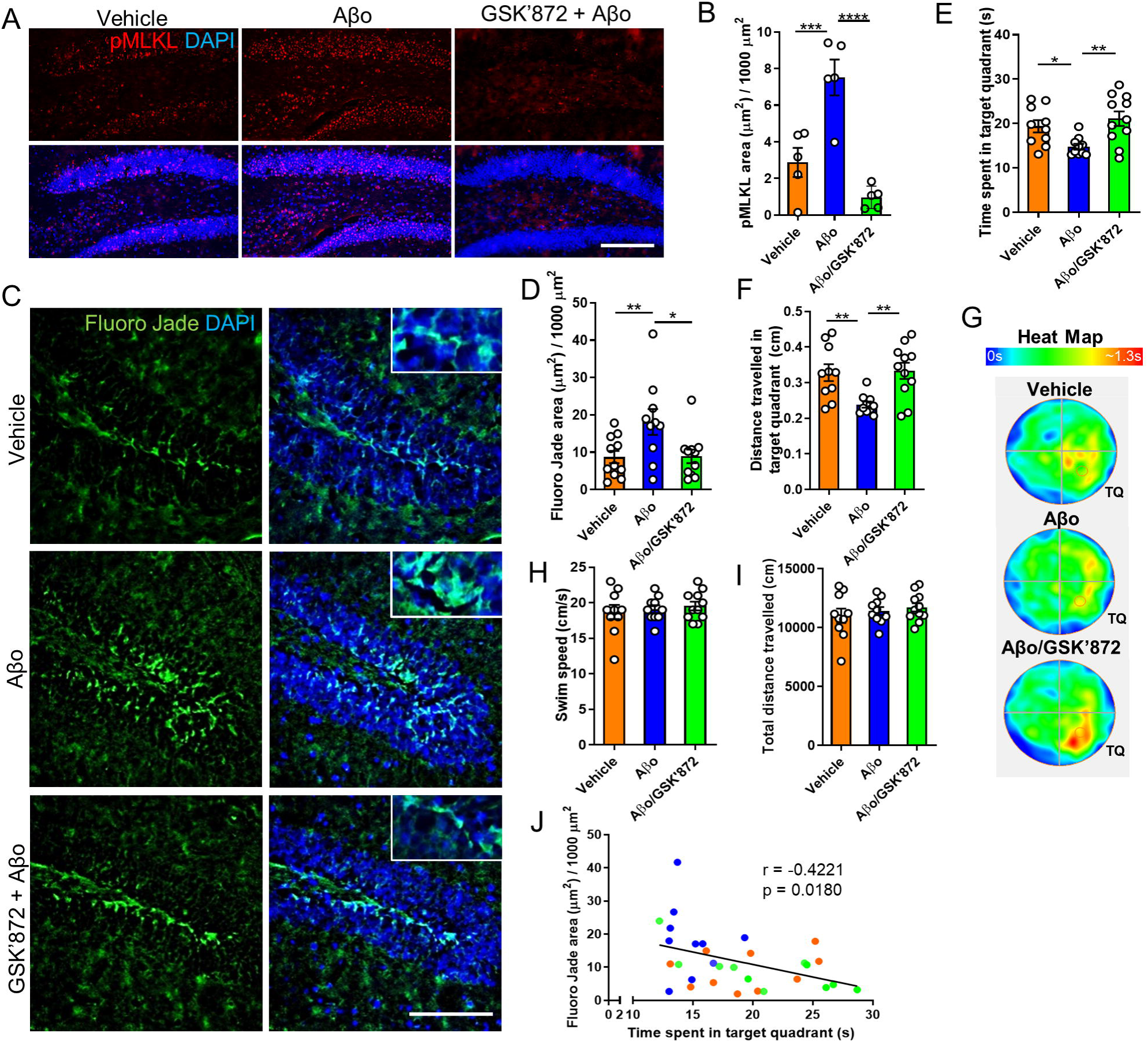
RIPK3 inhibition reduces neurodegeneration and memory loss induced by Aβo in mice. **(A**,**B)** Representative images and quantitative analysis of brain sections from wild-type mice treated as indicated in the images, immunolabeled with an anti-pMLKL antibody (scale bar, 150 μm). **(C**,**D)** Representative micrographs and quantitative measurement of brain sections from wild-type mice treated as indicated in the images, stained with Fluoro-Jade C (scale bar, 150 μm). n = 10 per group, 4 sections per brain, one image of the hippocampus per section was analyzed. **(E**,**F**,**H**,**I)** Graphs represent quantitative analyzes of the indicated parameters, obtained from the MWM test. **(G)** The heat map shows a graphic representation of the time that mice, treated as indicated in the image, spent in the target quadrant (TQ) during the MWM test. **(J)** Association analysis of Fluoro-Jade C stained area and cognitive performance was done by linear regression. Data are presented as mean ± S.E.M. Data in **B**,**D**-**F**,**H**, and **I** were analyzed by one-way ANOVA followed by Bonferroni post-test. Data in **J** were analyzed by linear regression using Pearson correlation. *p < 0.05; **p < 0.01; ***p < 0.001; ****p < 0.0001.

Together, these findings suggest that neurodegeneration induced by Aβo is mediated by the necroptotic machinery and that targeting necroptosis provides significant neuroprotection and results in cognitive improvement in experimental AD.

### Neurodegeneration triggered by Aβo-stimulated microglia occurs through necroptosis activation

As indicated above, AD brains are characterized by the presence of reactive microglial cells associated with Aβ aggregates (10–12, 14). This can be observed in human AD as well as in murine AD models (see Fig. 1C). Moreover, a role for microglia as mediators of Aβ-induced neurodegeneration has been extensively demonstrated (9). Interestingly, it has been reported that rather than fibrillar amyloid, soluble oligomeric Aβ structures can more effectively potentiate a proinflammatory microglial response leading to neurodegeneration (55–59). We then sought to explore whether microglial activation triggered by Aβo can elicit activation of the necroptotic pathway in neurons. Interestingly, *in vitro* experiments carried out with primary neuron-microglia co-cultures treated with Aβo revealed that Aβo-induced neurodegeneration is worsened when microglial cells are present (Supplementary Fig. 7). We used synthetic Aβo, and their cytotoxic properties were first assessed in primary neurons. Oligomers triggered progressive neurodegeneration, evidenced by beading and retraction of neuronal processes, that was extensive after 72 h of incubation (Supplementary Fig. 7A-C). Notably, the addition of microglial cells to neuronal cultures significantly enhanced Aβo neurotoxicity (Supplementary Fig. 7D,E).

To test the possible involvement of microglia-secreted factors to the induction of necroptosis-mediated neuronal death, hippocampal neurons were treated with conditioned medium from primary rat microglial cells stimulated with Aβo (CM^Aβ^, Fig. 5A). Treatment with CM^Aβ^ induced broad neuronal degeneration and the neurotoxic effects were dose-dependent (Fig. 5B,C). We then performed western blot analyses to determine the activation of necroptosis by analyzing the levels of RIPK3 and pMLKL in neurons upon treatment with CM^Aβ^. Densitometric analysis showed increased levels of RIPK3 (Fig 5D) and pMLKL (Fig 5E) in neurons exposed to CM^Aβ^ compared with neurons treated with CM^ctrl^ (full blots are shown in Supplementary Fig.8A,B). The physical interaction between RIPK1, RIPK3, and MLKL within the necrosome complex leads to MLKL phosphorylation, resulting in its oligomerization and execution of necroptosis (33). In order to further determine if CM^Aβ^ triggers the formation of the necrosome in neurons, co-immunoprecipitation experiments were performed followed by western blot analysis. These results revealed a significantly higher RIPK1-pMLKL interaction in neurons treated with CM^Aβ^ compared with neurons treated with CM^ctrl^, indicating that CM^Aβ^ was able to enhance the formation of the necrosome complex (Fig. 5F, full blots are shown in Supplementary Fig.8C). Finally, to define the involvement of necroptosis in neuronal death in this setting, we treated cells with the RIPK1 antagonist nec-1s or with GSK’872, prior to the addition of CM^Aβ^ to the cultures. Importantly, neurons treated with CM^Aβ^ together with nec-1s or with GSK’872 showed significant protection from neurodegeneration compared with neurons treated with CM^Aβ^ alone (Fig. 5G-J).

**Figure 5.**
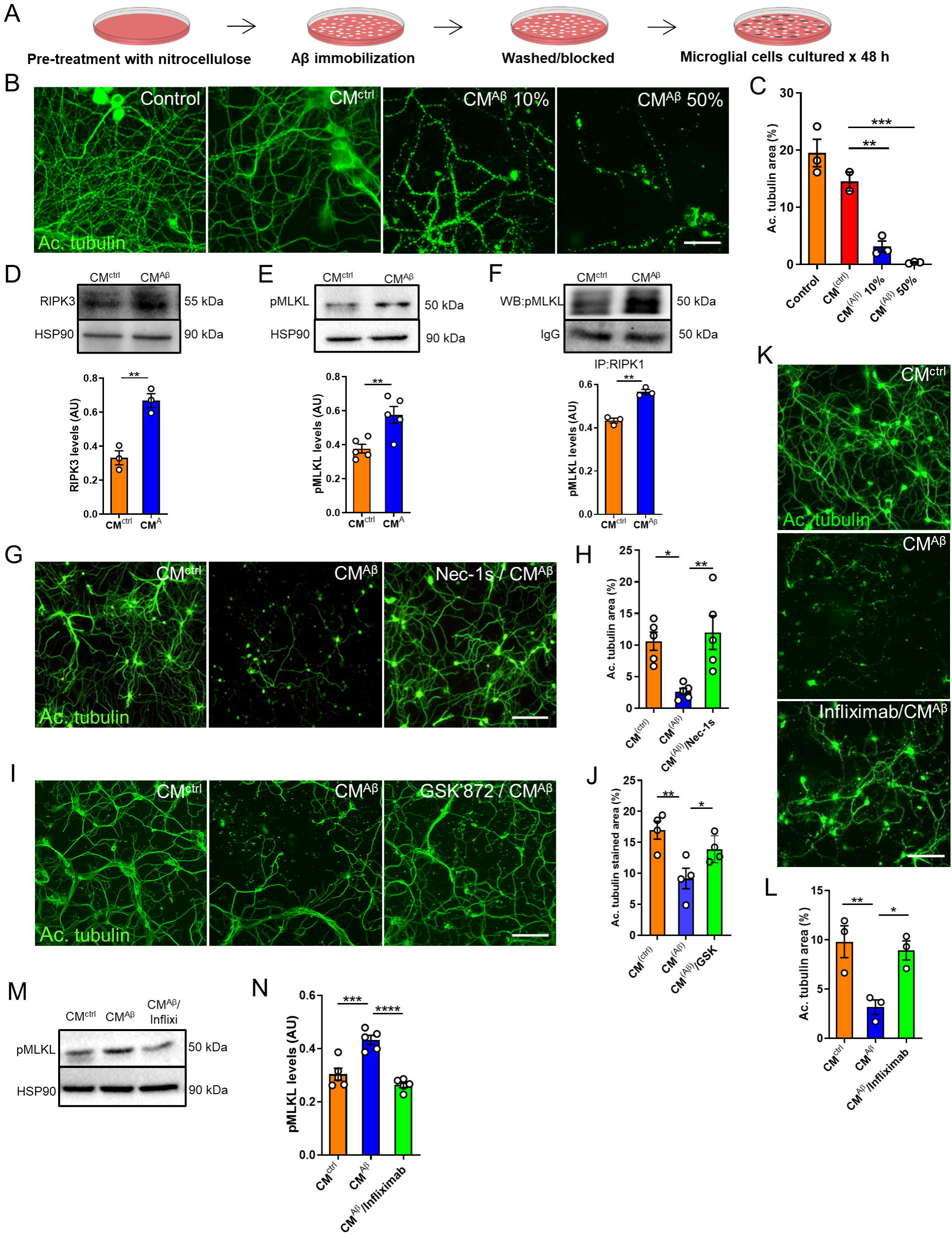
Neurodegeneration induced by conditioned medium from Aβo-stimulated microglia occurs via necroptosis activation. **(A)** Schematic representation of the preparation of the conditioned medium. **(B**,**C)** Representative images and quantitative analysis from neurons treated for 72 h with culture medium collected from a culture dish where Aβo were previously immobilized using nitrocellulose (control), neurons treated with conditioned medium from untreated microglia (CM^ctrl^), and neurons treated with 10% and 50% conditioned medium from Aβo-stimulated microglia (CM^Aβ^); immunostaining was performed with anti-acetylated tubulin antibody (scale bar, 50 μm). Western blot analyses of protein extracts from neurons treated for 6 h with CM were performed to determine the expression levels of RIPK3 **(D)** and pMLKL **(E)**. Specific bands were quantified by densitometric analysis and expressed relative to HSP90 protein, represented in the graphs. **(F)** Protein extracts from neurons treated for 6 h with CM were immunoprecipitated with an antibody against RIPK1 and probed for pMLKL. The graph represents the densitometric analysis of the western blots. **(G**,**H)** Representative micrographs and quantitative measurements of neurons treated for 72 h with CM^ctrl^, CM^Aβ^ and with Nec-1s (30 μM) prior to the addition of CM^Aβ^, immunolabeled with anti-acetylated tubulin antibody (scale bar, 200 μm). **(I**,**J)** Representative images and quantitative analysis of neurons treated for 72 h with CM^ctrl^, CM^Aβ^ and with GSK’872 (1 μM) prior to the addition of CM^Aβ^, immunolabeled with anti-acetylated tubulin antibody (scale bar, 200 μm). **(K**,**L)** Representative images and quantitative analysis of neurons treated with CM^ctrl^, CM^Aβ^, and with CM^Aβ^ that was previously mixed with Infliximab (0.01 μg/μl, Infliximab/CM^Aβ^), immunostained with anti-acetylated tubulin antibody (scale bar, 200 μm). **(M**,**N)** Western blot analysis of protein extracts from neurons treated for 6 h under the indicated conditions was performed to determine the expression levels of pMLKL. Bands were quantified by densitometric analysis and expressed relative to HSP90 protein, represented in the graph. Each experiment was performed at least three independent times, with three replicates per condition each time. Data are presented as mean ± S.E.M. Data in **C, H, J, L, and N** were analyzed by one-way ANOVA followed by Bonferroni post-test. Data in **D-F** were analyzed by Student’s *t-test*. *p < 0.05; **p < 0.01; ***p < 0.001, ****p < 0.0001.

Together, these data suggest that in response to stimulation with Aβo, neurotoxic factors released by activated microglia drive the formation of the necrosome and the initiation of the necroptosis signaling cascade in neurons.

### TNF-α released by Aβo-stimulated microglia contributes to necroptosis activation

The secretion of TNF-α by Aβo-stimulated microglial cells has been previously described (47). It has been demonstrated that activation of TNFR1 by TNF-α can trigger cell death via activation of the necroptotic pathway (31, 60). Hence, we next investigated the possible contribution of TNF-α to Aβo-induced necroptosis. CM^Aβ^ was incubated with the anti-TNF-α antibody Infliximab to neutralize the protein and was then used to treat neuronal cultures. Remarkably, Infliximab provided significant protection to neurons compared to neurons treated with control CM^Aβ^, which showed a high proportion of neuritic beading (Fig. 5K,L). To test whether the neuroprotection provided by Infliximab was due to the inhibition of necroptosis activation, we monitored the levels of pMLKL. Western blot analysis of neurons treated with CM^Aβ^ together with Infliximab showed decreased pMLKL levels compared to neurons treated with CM^Aβ^ alone (Fig. 5M,N; full blots are shown in Supplementary Fig.8D). To assess the sufficiency of TNF-α to trigger necroptosis-mediated neurodegeneration, we treated neuronal cultures with recombinant TNF-α. A concentration of 40 ng/ml was used, as it has been previously shown that when combined with apoptosis inhibitors, it is able to trigger necroptosis *in vitro* (61, 62). As shown in Supplementary Fig.9A,B, neither treatment with TNF-α alone nor in combination with CM^ctrl^ induced neurodegeneration.

Together, these data suggest that, although TNF-α is necessary, it acts in conjunction with other molecules released by Aβo-activated microglia to induce necroptosis-mediated neurodegeneration.

## Discussion

Several studies have demonstrated the involvement of necroptosis in the pathogenesis of neurodegenerative diseases, including PD (41), MS (40), ALS (39), Huntington disease (63), and ischemic neuronal injury (64). It has been recently shown that necroptosis is activated in human AD brains (26, 27). However, the mechanism by which this pathway is activated in AD is currently unknown. Here, we define a role for Aβ pathology in necroptosis activation in AD and demonstrate that microglial cells mediate Aβo-induced necroptosis activation in neurons.

The abnormal deposition of Aβ results from an imbalance between its production and clearance, constituting a crucial and probably causative event in the development of AD (2). Nevertheless, Aβ appears to interact with multiple receptors, different types of cells and affect many biological processes and still, the exact mechanism by which neurons die in AD is not clear. Our results demonstrate a spatial correlation between Aβ pathology and the necroptosis effector pMLKL in AD brains and a correlation between Aβo burden and the expression of key necroptosis drivers, suggesting a possible link between Aβ aggregation and the activation of the necroptosis signaling pathway. Whereas Aβ plaques have long been a signature of AD histopathology, increasing studies over the past decade have demonstrated that soluble oligomeric Aβ constitutes the most neurotoxic type of Aβ aggregate (3, 5, 65, 66). Accordingly, Brody *et al*., hypothesized that plaques may serve as a reservoir for toxic oligomers, thereby capturing them in the extracellular space. It was suggested that over time, plaques get saturated and therefore oligomers can diffuse and exert their toxicity to the surrounding cells (4). In agreement with this hypothesis, Koffie and colleagues demonstrated that oligomeric Aβ associated with plaques colocalizes with postsynaptic densities and is associated with spine collapse and synapse loss in postmortem AD samples (7). Consistent with this evidence, our data point to oligomeric Aβ as a driver of necroptosis activation in AD, leading to neurodegeneration and memory loss. Although a previous report found no correlation between Aβ and necroptosis markers in AD brains, in this study the researchers considered Aβ plaque load and not oligomers for the analysis (26). Many studies have demonstrated that the aggregation process of Aβ follows a seeding-nucleation mechanism that involves the formation of multiple types of intermediates including oligomers and fibrils (67, 68). Additional studies are necessary to test the role of insoluble fibrillar Aβ in the activation of necroptosis. In addition, previous studies have shown colocalization between pTau and necroptosis markers in AD brains (26, 27), which is not surprising considering the mounting evidence proving the synergistic contribution of both misfolded Aβ and tau proteins to neurodegeneration (61–64). Further studies would be required in order to test a potential interplay between both proteins for necroptosis activation.

In agreement with the study by Caccamo *et al*. (26), our results suggest that inhibition of the necroptotic pathway prevents neurodegeneration in AD. Previous data from our group provided evidence for a functional role of necroptosis in PD, as necroptosis inhibition improved motor performance in a PD model (41). Importantly, the present study is the first to unveil the functional implications of targeting necroptosis in an experimental AD model. As necroptosis is a druggable pathogenic mechanism, our findings suggest that necroptosis inhibition has an important translational potential for AD.

Several pathological processes may trigger activation of the necroptotic cascade in AD, including inflammation and oxidative stress. Our data suggest that upon interaction with Aβo, microglial cells secrete cytotoxic molecules that induce neurodegeneration via necroptosis activation. Among the secretory mediators released by microglia, TNF-α elicits necroptosis *in vitro* and *in vivo* (31, 60). Accordingly, previous studies in other neurodegenerative conditions, including MS and ALS, demonstrated that sustained inflammation is associated with TNF-α-mediated necroptosis (73, 74). In line with this evidence, our work suggests that TNF-α released from microglia following its activation by Aβo contributes to necroptosis-mediated neurodegeneration. Aβ can stimulate signaling responses in microglial cells that induce the release of a number of molecules that can contribute to neurodegeneration. Interestingly, there is evidence showing that glutamate, which has been previously demonstrated to be released by activated microglia (75, 76), can also induce necroptosis activation (77, 78). Similarly, interferon-γ, which is produced by reactive microglia (79), has also been shown to participate in the activation of the necroptotic signaling pathway (29). Additional studies are needed to unveil the microglial protein expression profile in response to Aβo stimulation and define the exact molecules involved in the activation of the necroptotic pathway.

Overall, in the present study, we provide new insights into the mechanism by which necroptosis is activated in AD, adding a new layer of complexity involving the crosstalk between microglia and neurons. Our results may contribute to understanding the mechanisms that lead to neurodegeneration in AD and support the view of necroptosis as a potential therapeutic target for disease intervention.

## Acknowledgments

We thank Dra. Claudia Duran for suggestions and Dr. Claudio Hetz for critical review of the manuscript.

## Funding

This study was funded by grants from FONDECYT No. 3180341 to NS; FONDECYT No. 1150766 and FONDAP program 15150012 to FC; Instituto de Salud Carlos III (ISCiii) of Spain, co-financed by FEDER funds from European Union, through grants PI18/01556, CIBERNED (CB06/05/0094) and by Junta de Andalucia Consejería de Economía y Conocimiento (US-1262734) to JV; Ministry of Science and Innovation (MICIN) State Research Agency grants PID2019-107090RA-I00 and Ramon y Cajal Program RYC-2017-21879 to IMG and grants from NIH R01AG059321 and R01AG061069 to CS.

The authors declare no competing financial interests.

The authors confirm that the data supporting the findings of this study are available within the article and its supplementary materials.

## Author Contributions

N.S. designed research; N.S., I.M., N.G., G.Q., L.V., S.J., and M.E. performed research; N.S. and J.V. analyzed data; C.S., A.G., and J.V. contributed new reagents or analytic tools; N.S. wrote the paper; F.C. supervised research and edited the paper.

## Figure Legends

**Supplementary Figure 1. Validation of the pMLKL immunoreactivity by DAB staining**. Representative micrographs of hippocampal brain areas of AD patients (n = 3) immunostained with anti-pMLKL antibody. Orange arrows indicate pMLKL-positive neurons, and green arrows show pMLKL-positive microglia. Magnification, 20X.

**Supplementary Figure 2. Coefficient of variation of the dot-blot assays**. The coefficient of variation (CV) was calculated using triplicates of Braak V-VI samples. A CV of 11.63% and 15.06% was determined for OC and A11, respectively.

**Supplementary Figure 3. Validation of the pMLKL antibody and assessment of cell specificity**. Representative micrographs of **(A)** the dentate gyrus (DG) and **(B)** CA1 brain regions from wild-type and MLKL knockout mice subjected to intracerebral injection of Aβo, immunostained with anti-pMLKL antibody (scale bar, DG: 100 μm, CA1: 150 μm). **(C**,**D)** Representative images of the hilus of wild-type mice treated with Aβo, labeled with the indicated antibodies (scale bar, 50 μm and 25 μm for magnifications).

**Supplementary Figure 4. Intracerebral administration of Aβo elicits neurodegeneration in wild-type mice**. Representative micrographs of brain sections from wild-type mice treated as indicated in the images, stained with Fluoro-Jade C (scale bar, 150 μm).

**Supplementary Figure 5. MLKL inhibition attenuates Aβo-induced microgliosis in mice. (A**,**B)** Representative micrographs and quantitative analysis of brain sections from wild-type and *Mlkl* knockout mice treated as indicated in the images, immunostained with anti-Iba1 antibody (scale bar, 200 μm). The data are presented as mean ± S.E.M. and were analyzed by one-way ANOVA followed by Bonferroni post-test. *p < 0.05.

**Supplementary Figure 6. RIPK3 inhibition does not alter Aβo-induced microgliosis in wild-type mice. (A**,**B)** Representative micrographs and quantitative analysis of brain sections from wild-type mice treated as indicated in the images, immunostained with anti-Iba1 antibody (scale bar, 200 μm). The data are presented as mean ± S.E.M. and were analyzed by one-way ANOVA followed by Bonferroni post-test. *p < 0.05; **p < 0.01.

**Supplementary Figure 7. Aβo neurotoxicity is worsened when microglial cells are present. (A)** To characterize Aβo, a negative stain with 1% uranyl acetate was performed followed by electron microscopy analysis. Arrows indicate small oligomeric structures (scale bar, 200 nm). **(B**,**C)** Representative images and quantitative analysis of neurons treated as indicated, immunostained with anti-acetylated tubulin antibody (scale bar, 100 μm). **(D**,**E)** Representative micrographs and quantitative measurement of neurons and neuron/microglia cocultures treated as indicated, immunolabeled with anti-acetylated tubulin and anti-Iba1 antibodies as specified in the images (scale bar, 100 μm in the middle panel; 50 μm in the right panel). Each experiment was performed at least three independent times, with three replicates per condition each time. The data are presented as mean ± S.E.M. and were analyzed by one-way ANOVA followed by Bonferroni post-test. *p < 0.05; **p < 0.01; ****p < 0.0001

**Supplementary Figure 8. Full blots of pMLKL and RIPK3 western blots. (A)** The membrane was first probed with an anti-hsp90 antibody and then incubated with anti-RIPK3. **(B)** The membrane was cut, and each piece was incubated with the indicated antibody. Left panel: colorimetric image showing protein ladder. **(C)** The membrane was cut and incubated with an anti-pMLKL antibody. Following stripping, and after confirming that no bands could be visualized with ECL, membranes were blocked, and detection of the IgG was done. Left panel: colorimetric image showing protein ladder. **(D)** The membrane was cut, and each piece was incubated with the indicated antibody. Left panel: colorimetric image showing protein ladder.

**Supplementary Figure 9. TNF-α alone is not sufficient to trigger neuronal necroptosis. (G**,**H)** Representative micrographs and quantitative measurements of neurons treated for 72 h with CM^ctrl^, CM^Aβ^, TNF-α, and with CM^ctrl^/TNF-α, immunolabeled with anti-acetylated tubulin antibody (scale bar, 200 μm). The data are presented as mean ± S.E.M. and were analyzed by one-way ANOVA followed by Bonferroni post-test. *p < 0.05; **p < 0.01.

