## Supplementary figures and images for "Aβ oligomers trigger necroptosis-mediated neurodegeneration via microglia activation in Alzheimer’s disease"

### Supplemental Figure 1

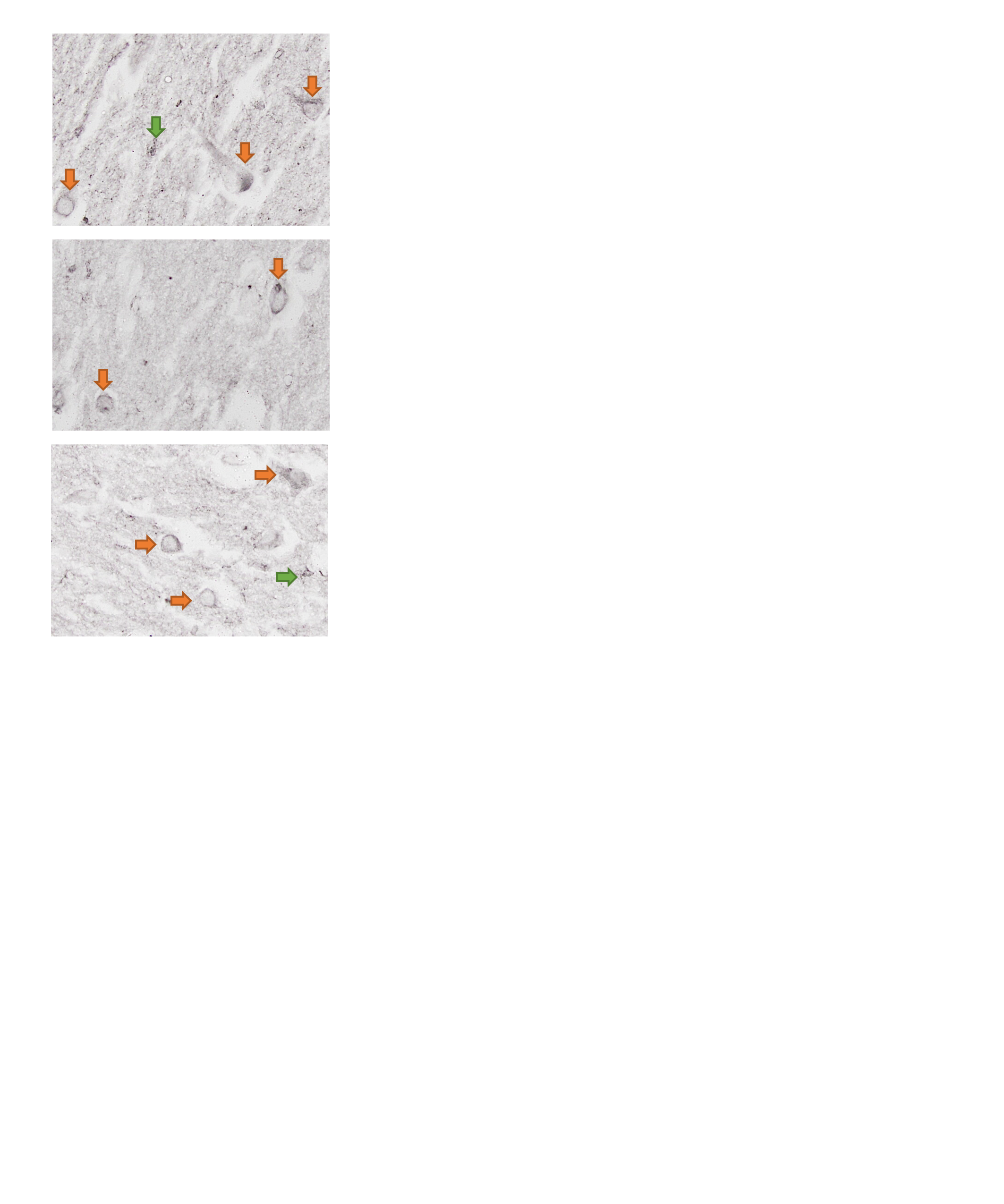

### Supplemental Figure 2

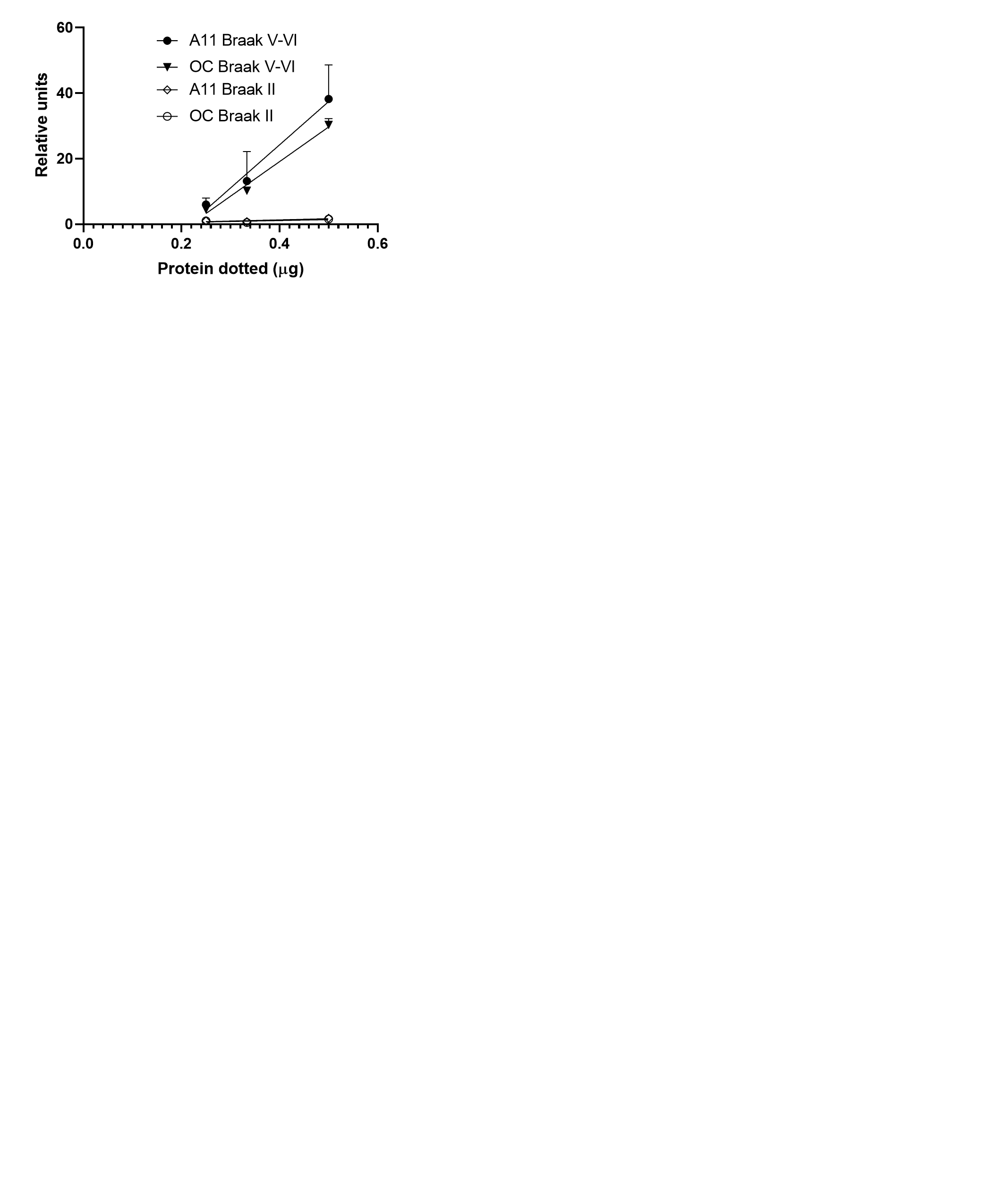

### Supplemental Figure 3

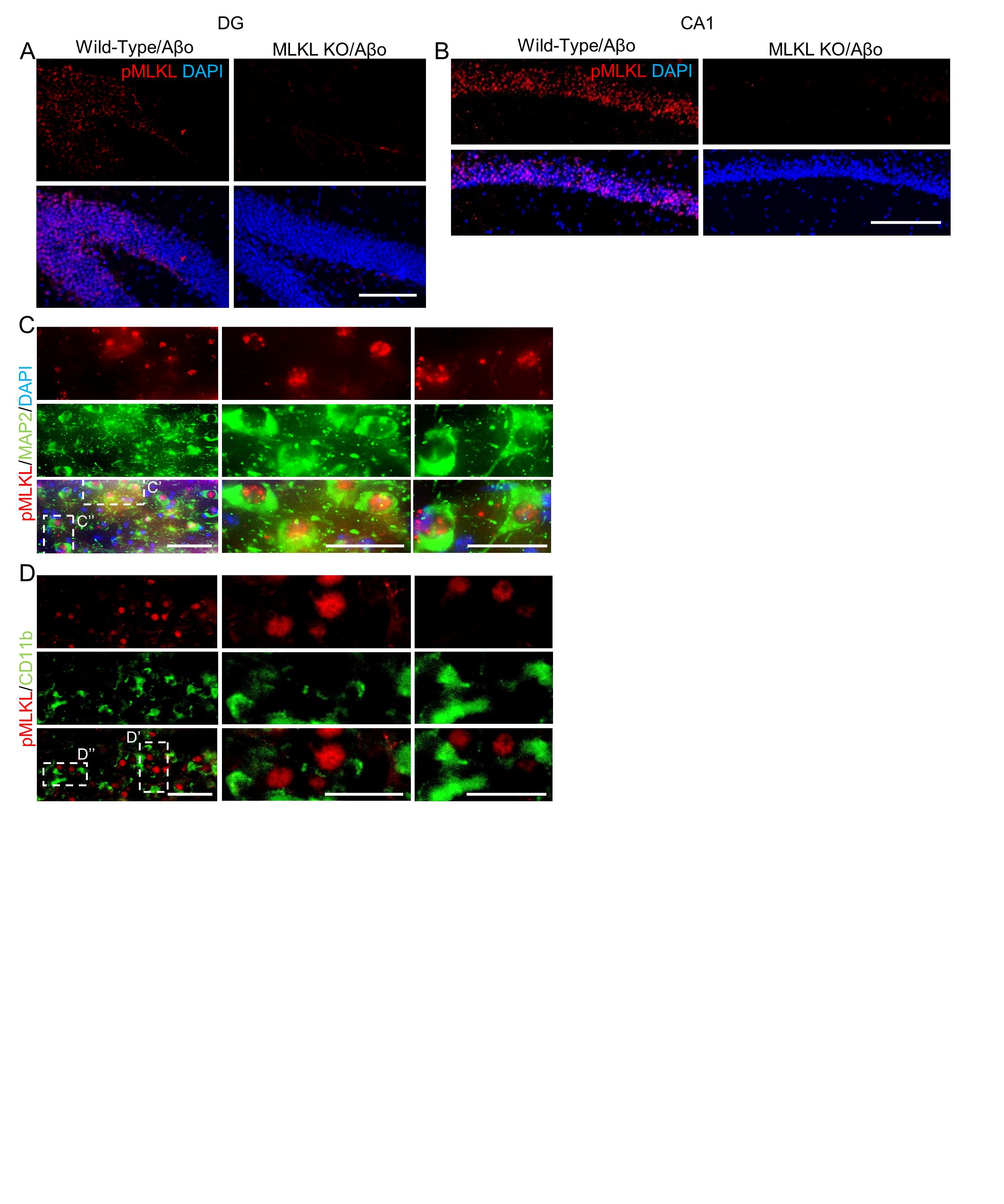

### Supplemental Figure 4

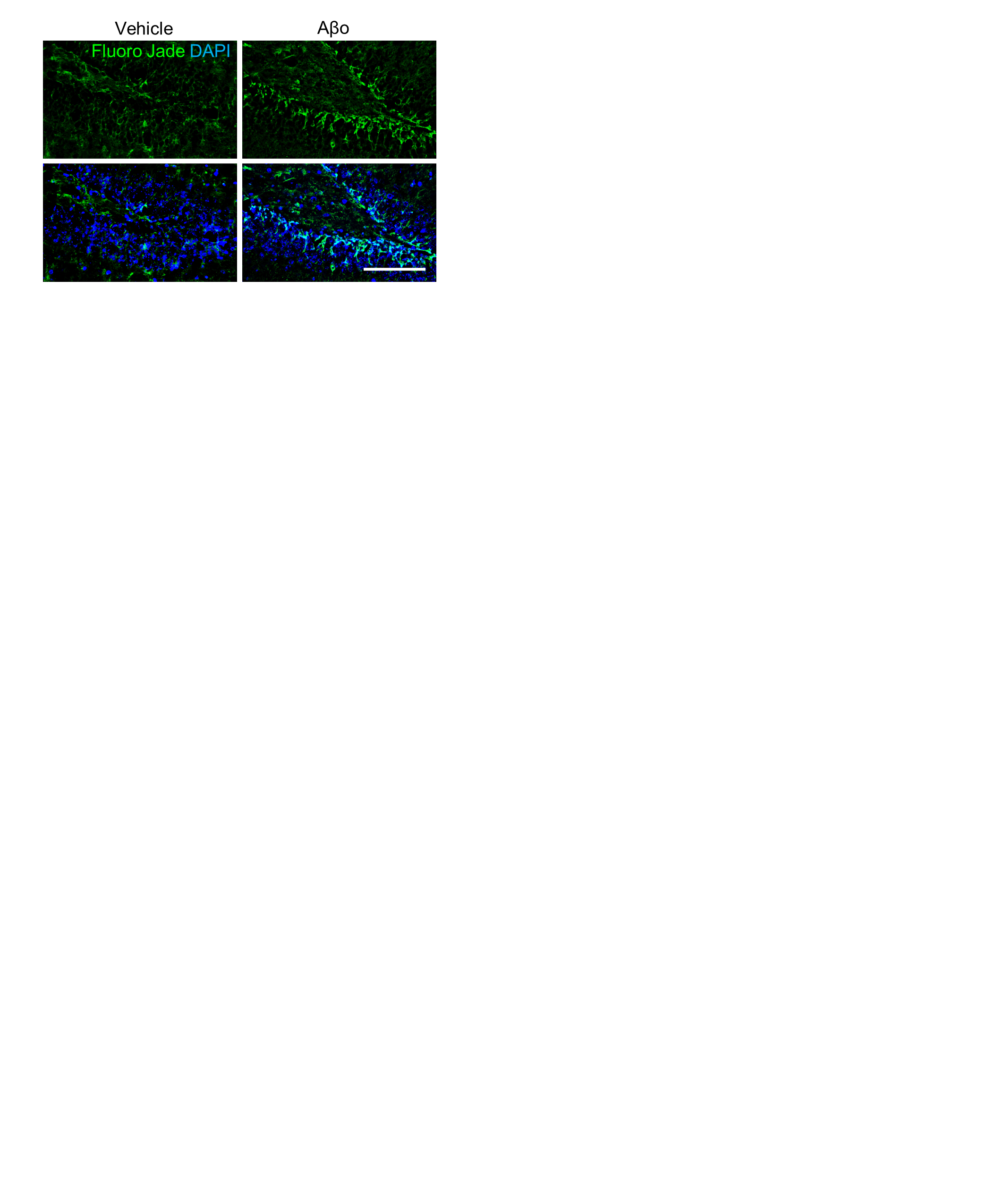

### Supplemental Figure 5

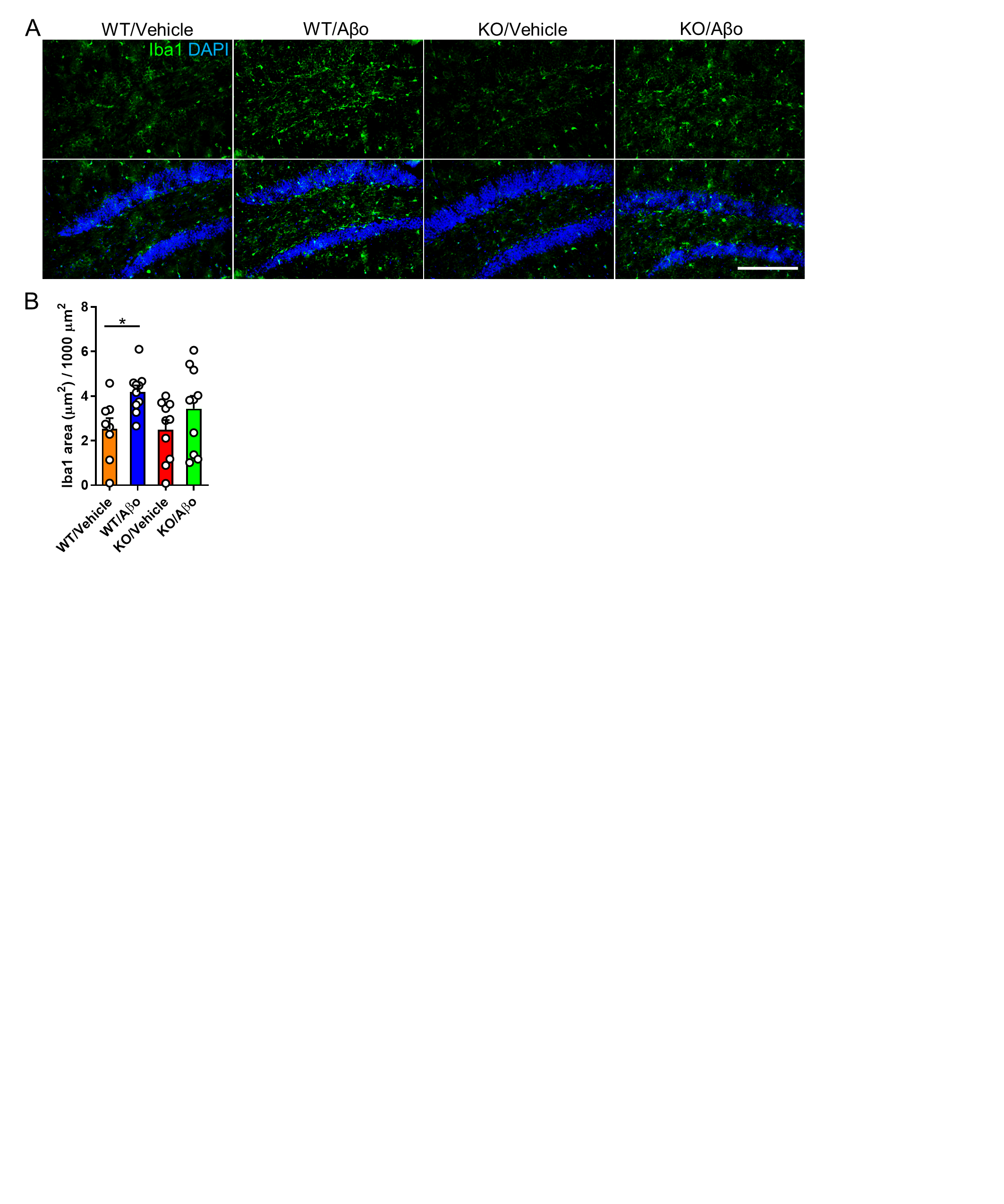

### Supplemental Figure 6

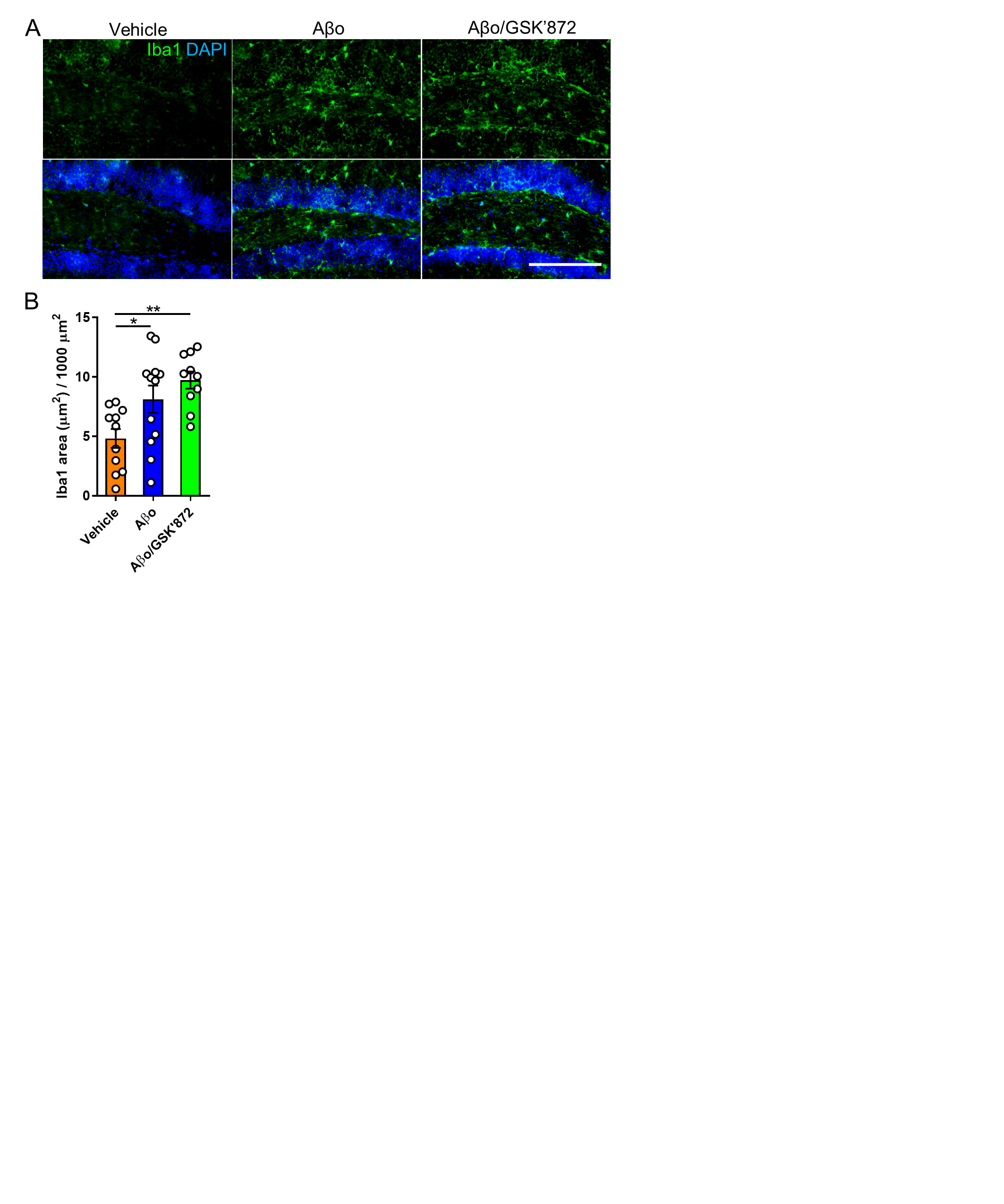

### Supplemental Figure 7

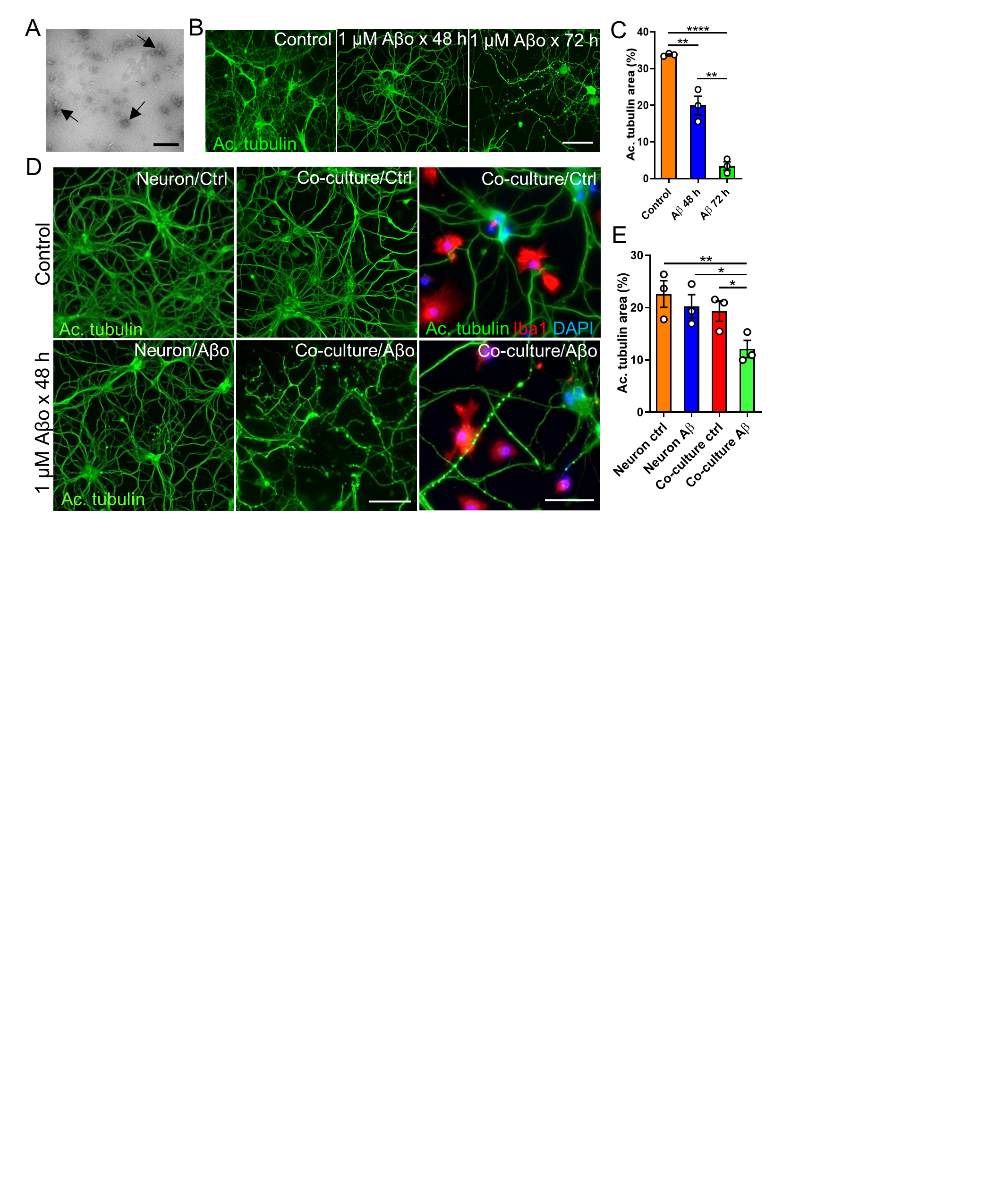

### Supplemental Figure 8

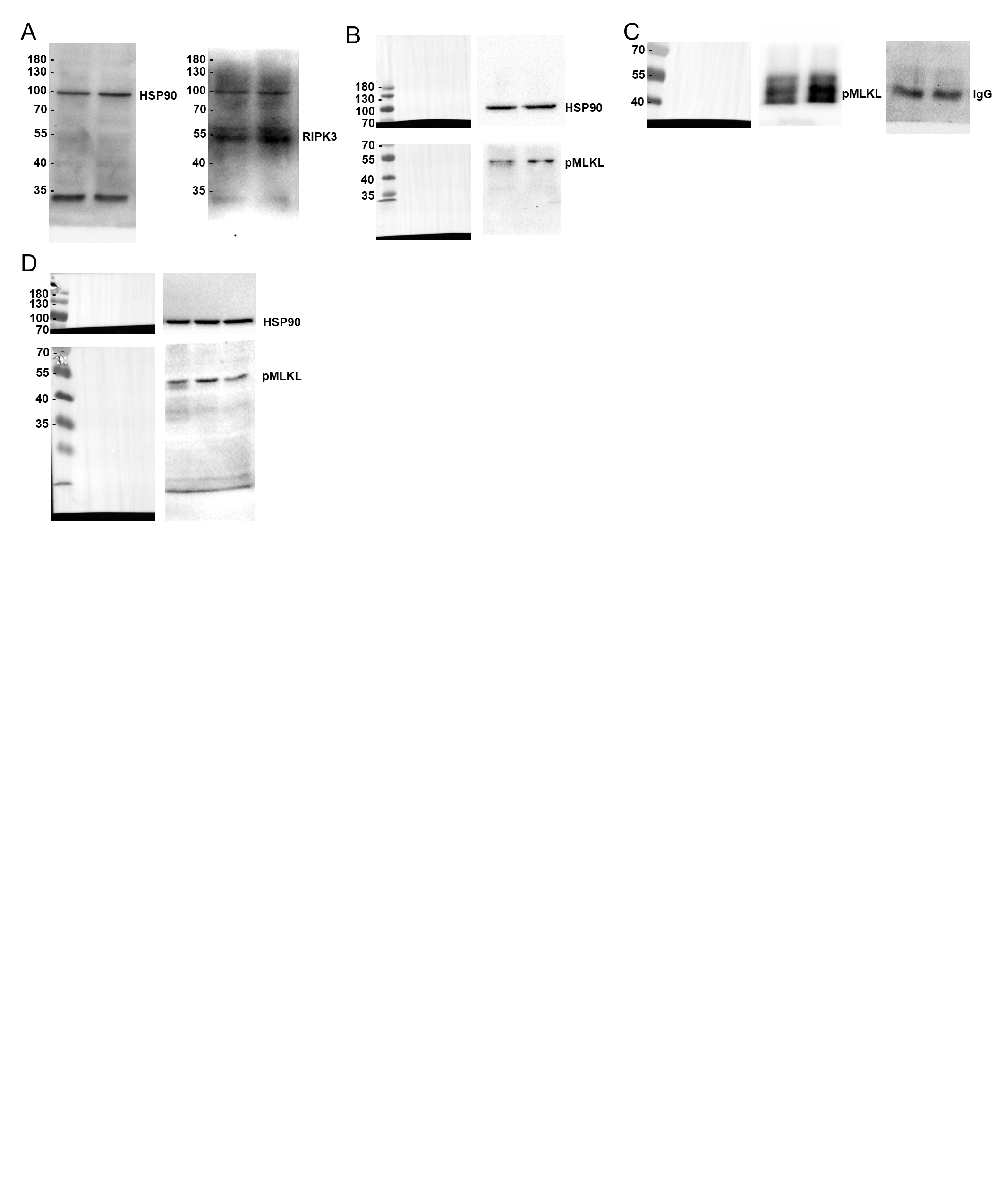

### Supplemental Figure 9

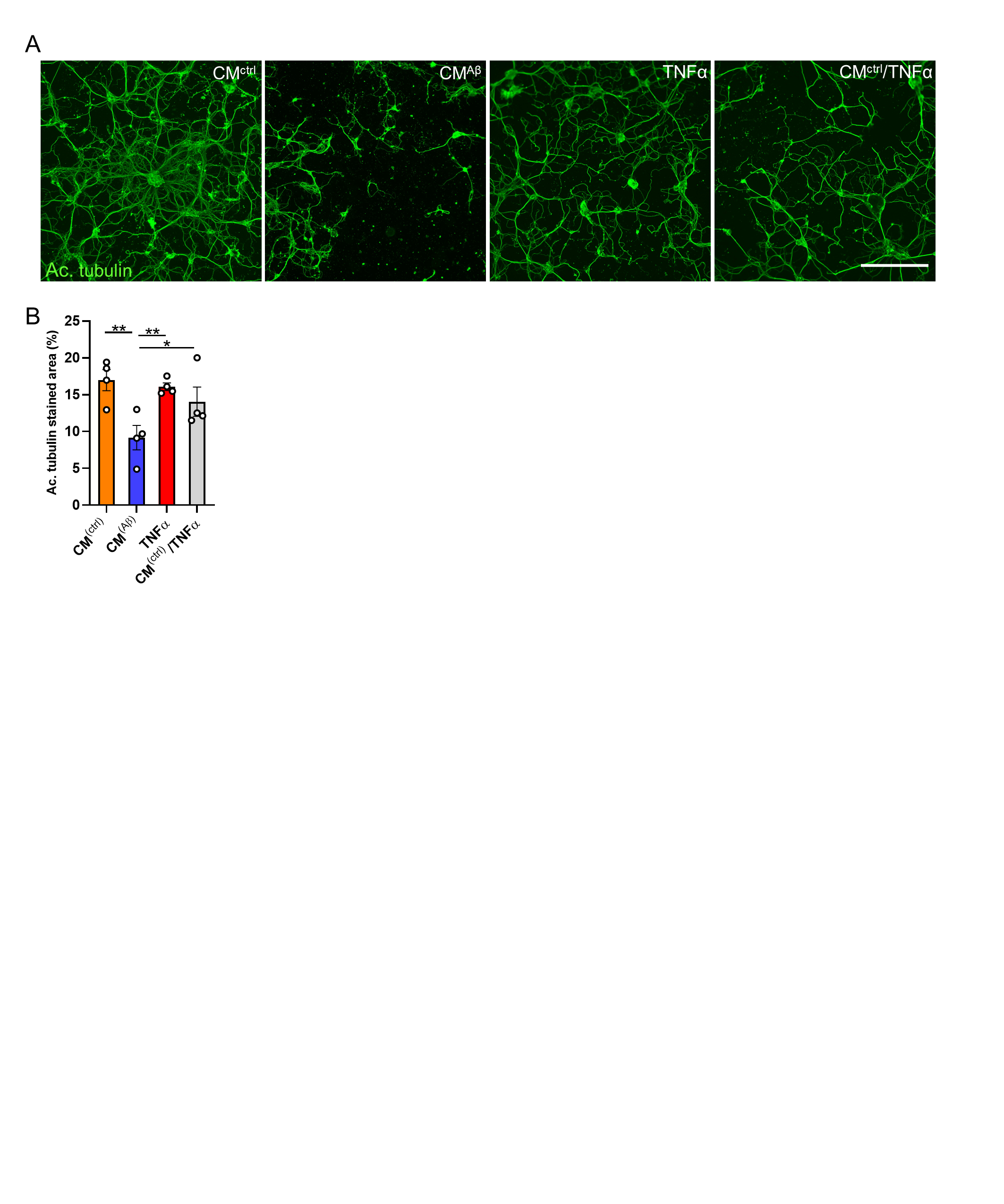
